# Platelet, erythrocyte, endothelial, and monocyte microparticles in coagulation activation and propagation

**DOI:** 10.1101/2020.01.06.895722

**Authors:** E.N. Lipets, O.A. Antonova, O.N. Shustova, K.V. Losenkova, A.V. Mazurov, F.I. Ataullakhanov

## Abstract

**Background and objective:** For many pathological states, microparticles are supposed to be one of the causes of hypercoagulation. Although there are some indirect data about microparticles participation in coagulation activation and propagation, the integral hemostasis test Thrombodynamics allows to measure micropaticles participation in these two coagulation phases directly by influence on the appearance of coagulation centers in plasma volume and on the rate of clot grown from surface with immobilized tissue factor.

**Methods:** Microparticles were obtained from platelets and erythrocytes by stimulation with SFLLRN and A23187, respectively, from monocytes, endothelial HUVEC culture and monocytic THP cell culture by stimulation with lipopolysaccharides. Microparticles were counted by flow cytometry and titrated in microparticle-depleted normal plasma in the Thrombodynamics test.

**Results:** Monocyte microparticles induced the appearance of clotting centres through the TF pathway at concentrations approximately 100-fold lower than platelet and erythrocyte microparticles, which activated plasma by the contact pathway. For endothelial microparticles, both activation pathways were essential, and their activity was intermediate. Monocyte microparticles induced plasma clotting by the appearance of hundreds of clots with an extremely slow growth rate, while erythrocyte microparticles induced the appearance of a few clots with a growth rate similar to that from surface covered with high-density tissue factor. Patterns of clotting induced by platelet and endothelial microparticles were intermediate. Platelet, erythrocyte and endothelial microparticles impacts on the rate of clot growth from the surface with tissue factor did not differ significantly within the 0-200·10^3^/ul range of microparticles concentrations. However, at concentrations greater than 500·10^3^/ul, erythrocyte microparticles increased the stationary clot growth rate to significantly higher levels than do platelet microparticles or artificial phospholipid vesicles consisting of phosphatidylcholine and phosphatidylserine.

**Conclusion:** Microparticles of different origins demonstrated qualitatively different characteristics related to coagulation activation and propagation.

## Introduction

Cell destruction or activation leads to microparticles (MPs) shedding. In the blood of normal donors, more than 80% of MPs are derived from platelets [1, 2]. In pathological states the MPs concentration and origin may change.

There are a number of clinical works where the MPs concentration is shown to increase in pathological states associated with elevated thrombotic risk. Data are represented in corresponding reviews [3–8]. Many studies have also revealed elevated MPs concentration in both arterial and venous thrombosis. In acute coronary syndromes, the concentration of platelet MPs (PMPs) was found to be increased [9], as was the concentration of endothelial MPs (EMPs) [10–12]. Increased tissue factor- bearing MPs concentration [13, 14] or activity [15, 16] was associated with venous thromboembolic events in patients with cancer. At the same time, data about the MPs concentration in unprovoked venous thromboembolism (VTE) are very contradictive [5, 17]. Recently, a number of prospective studies have appeared that examine the role of increased concentrations of MPs in cancer and recurrent thrombosis, but their results are also contradictory [5, 17]. For other pathological states, prospective studies are few: EMPs concentration increase was an independent predictor of cardiovascular complications in patients with heart failure, type II diabetes, and end-stage renal failure [18–20].

Thus, although there is a considerable amount of data on the involvement of MPs in hypercoagulation, MPs concentration does not always reflect the risk of thrombotic complications. Therefore, the study of mechanisms of MPs influence on coagulation is of current interest. It is well known that coagulation can be activated by two pathways: external, when coagulation is triggered by tissue factor (TF), and internal, when factor XII is activated. This phase is called coagulation initiation. Activation of either pathway leads to the same reactions in the coagulation cascade – the clot growth phase. The process finishes with the termination phase – the protein C reactions. Phospholipid surfaces containing phosphatidylserine (PS) play an important role in these phases, especially in the propagation phase. MPs are able to increase the clot growth rate by providing such surfaces and by binding coagulation factors at higher concentrations than platelets [21–23]. MPs can also trigger coagulation activation. TF-dependent activation from monocyte MPs (MMPs), obtained by monocyte stimulation with lipopolysaccharide (LPS), and a weaker contact activation from PMPs were shown in the works [24–26] in the thrombin generation test without the addition of activator. Contact activation from erythrocyte MPs (ErMPs) was shown in the works of van der Meijden [26] and Rubin et al. [27]. The stronger activation from TF-bearing MMPs and EMPs compared to TF-negative PMPs, ErMPs, and neutrophil MPs was demonstrated in the recalcification test in works [28–30]. The importance of PS in the procoagulant effect of PMPs [25] and the amplification of TF-initiated thrombin generation by PMPs [24] and ErMPs [27] indirectly indicate their participation in coagulation propagation. However, all these works were performed with homogeneous tests that do not allow for direct separation of the activation and propagation phases.

Thrombodynamics test allows us to observe the activation and propagation phases independently. The time of appearance and number of spontaneous activation centres characterize the ability of MPs to initiate coagulation. The MPs influence on the growth rate of clot triggered by surface with immobilized TF characterizes their impact on coagulation propagation. In this work, we compared MPs derived from almost all cells that are the major sources of MPs in blood (platelets, erythrocytes, endothelial cells and monocytes) in Thrombodynamics test. The comparison revealed the predominant impact of TF-bearing MMPs and EMPs on coagulation activation and ErMPs on coagulation propagation.

## Materials and methods

### Preparation of microparticles of different cellular origins

Microparticles (MPs) produced by activated platelets, monocytes, human monocytic THP-1 cells and endothelial cells (ECs) were prepared as described earlier in detail [29, 31]. Platelets and monocytes were isolated from the blood of healthy volunteers. The study was approved by the ethics committee of Center for Theoretical Problems of Physicochemical Pharmacology and and written informed consent was obtained from all donors. Washed platelets suspended at a concentration of 5 x 10^8^ platelets/ml in Tyrode/HEPES solution (137 mM NaCl, 2.7 mM KCl, 0.36 mM NaH_2_PO_4_, 0.1% dextrose, 1 mM MgCl_2_, 1 mM CaCl_2_, 0.35% BSA, 5 mM HEPES, pH 7.35) were activated by 10 µM thrombin receptor activating peptide (SFLLRN) for 10 min at 37°C.

For the isolation of monocytes, blood was centrifuged at 180 *g* for 10 min to obtain platelet- rich plasma, platelets were removed by centrifugation (1000 *g*, 15 min), and plasma was returned to the blood. Starch solution was added to agglutinate erythrocytes, and after their precipitation, the supernatant was layered over Histopaque1077 solution and centrifuged at 400 *g* for 30 min. Mononuclear leukocytes were washed from the “buffy coat” formed after this centrifugation; resuspended in RPMI 1640 medium containing 20 mM HEPES, 2 mM L-glutamine, 1 mM sodium pyruvate, penicillin (50 U/ml), streptomycin (100 μg/ml), and 10% fetal bovine serum; seeded into untreated 100-mm Petri dishes (25 x 10^6^ cells/dish); and cultivated for 18 hours. Non-attached mononuclear cells were washed off, and adherent monocytes were activated by 1 µg/ml bacterial lipopolysaccharide (LPS) for 6 hours. Activation of monocytes by LPS (as well as activation of ECs and THP-1 cells – see below) was performed in a medium containing non-inactivated fetal bovine serum (instead of inactivated) as a source of LPS-binding protein. THP-1 cells were obtained from the American Type Culture Collection (ATCC, Bethesda, MD), cultured under standard conditions (RPMI 1640 medium, containing 20 mM HEPES, 2 mM L-glutamine, 1 mM sodium pyruvate, penicillin (50 U/ml), streptomycin (100 μg/ml), and 10% fetal bovine serum), and activated by 1 µg/ml LPS for 6 hours.

ECs obtained from the human umbilical vein were cultured under standard conditions (DMEM medium containing 20 mM HEPES, 2 mM L glutamine, 1 mM sodium pyruvate, penicillin (50 U/ml), streptomycin (100 μg/ml), 10% heat inactivated fetal bovine serum, 200 μg/ml vascular endothelial growth factor, and 5 U/ml heparin) and activated by 1 µg/ml LPS for 12 hours.

Erythrocyte-derived MPs were prepared as described by Van Der Meijden et al. [32] with some modifications [30]. Blood from healthy volunteers was collected in 3.8% sodium citrate at a blood/anticoagulant ratio of 9/1. Blood was centrifuged at 180 *g* for 10 min. Platelet-rich plasma and the leukocyte “buffy coat” were removed, and 1 ml of erythrocytes was collected from the lower part of the erythrocyte pellet and diluted in 9 ml of HBS buffer. Erythrocytes were counted in an Abacus Junior B haematological analyser (Diatron Ltd., Austria) and washed 3 times in HBS at 2000 *g* for 15 min. Erythrocytes were resuspended in HBS at a concentration of 1 x 10^9^/ml, supplemented with 3 mM CaCl_2_ and treated with 10 µM A23187 calcium ionophore (Sigma-Aldrich, Inc., St. Louis, MI, USA) for 60 min at room temperature.

Activated platelets were spun down by double centrifugation at 2500 *g* for 15 min. Culture medium from activated monocytes and ECs and suspension of activated THP-1 cells were centrifuged at 400 *g* for 10 min and then at 2500 *g* for 15 min. Erythrocytes and their large fragments were removed by centrifugation at 2000 *g* for 15 min and at 2500 *g*. MPs were sedimented from obtained supernatants at 20,000 *g* for 30 min at 4°C and resuspended in filtered (filters Millex® - VV, 0.1 µm) HEPES- buffered saline (HBS, 10 mM HEPES, 140 mM NaCl, pH 7.4) containing 1% BSA (HBS/BSA). Suspensions contained in 1 ml MPs from 5 x 10^8^ platelets, 1 x 10^6^ monocytes, 1 or 3 x 10^6^ THP-1 cells, 1 x 10^6^ ECs, and 1 x 10^9^ erythrocytes.

All MPs were frozen in liquid nitrogen, stored at -70°C for no longer than 6 months and thawed at 37°C just before use. Repeat freezing/thawing cycles were avoided. For procoagulant activity measurement, MPs were thawed, centrifuged for 30 min at 16,000 *g*, resuspended in buffer A (150 mM NaCl, 2.7 mM KCl, 1 mM MgCl2, 0.4 mM NaH2PO4, 20 mM HEPES, 5 mM glucose, 0.5% bovine serum albumin, pH 7.4, filtered through a 0.22-µm membrane) and concentrated to the necessary extent.

### Counting of microparticles

MPs were thawed at 37°C, and 5 to 45 µl of suspension was added to 300 µl of annexin V binding buffer (Becton Dickinson, BD Bioscience, San Jose, CA, USA). After the addition of 2.5 µl of annexin V-FITC (Becton Dickinson, BD Bioscience, San Jose, CA, USA), the suspension was incubated in the dark for 30 min at room temperature. Control probes contained filtered HBS buffer without MPs. Standard beads with a diameter of 3 µm and known concentration (MP Count Beads, BioCytex, Stago, France) were used for counting calibration. Fifteen microliters of these beads was added to the analysed probes. Standard beads with a diameter of 1 µm (Flow cytometry Sub-Micron Size Reference Kit, Invitrogen, Life Technologies Corp., Carlsbad, CA, USA) were used for sizing calibration. MPs were analysed and counted in a FACS Canto II flow cytometer (Becton Dickinson, BD Bioscience, San Jose, CA, USA). Events were acquired at a low rate mode (less than 3000 events per sec). Data acquisition and analysis were performed using CELL Quest TM software (Becton Dickinson, BD Biosciences, San Jose, CA, USA). The noise threshold was set up in the FITC fluorescence channel (200 arbitrary units, a.u.). Events above this threshold were gated and counted in the SSC/FSC (side scattering/forward scattering) window in the size gate < 1 µm (size calibration beads). Analysis was performed until 1000 events were acquired in the gate for 3-µm counting beads. Examples of MPs analysis and counting are presented in Fig. 1. The concentration of MPs in probes was calculated as follows: number of MPs in 1 µl = (number of events in the gate < 1 µm) х (number of counting beads in 1 µl) x (dilution coefficient) / 1000. The dilution coefficient was applied upon the addition of different volumes of analysed MPs (from 5 to 45 µl) and counting beads (15 µl). The number of events (< 1 µm) in 1 µl of negative control probes (HBS buffer without MPs) was subtracted from the number of MPs in 1 µl. For all types of MPs, we observed a linear relationship between the number of counted MPs and the volume of analysed probe (5-45 µl). In preliminary experiments, we noticed that a much lower number of MPs were counted when the noise threshold was set up in the SSC or FSC channel, which was presumably due to the elimination of a significant fraction of small MPs from the analysis. We also noticed that the percentage of annexin V-positive MPs (above the threshold of the negative control) significantly varied in MPs of different cellular origins (for examples, see Fig. 1). The smallest percentage was detected for MPs from erythrocytes (lower than 10%), intermediate percentages for MPs from platelets (15-20%), higher percentages for MPs from monocytes and THP-1 cells (approximately 30%), and the highest percentage for MPs from ECs (up to 40%). Because of these variations, we counted not only annexin V-positive but all MPs and used these counts in comparative studies of their coagulation activity.

**Fig. 1.**
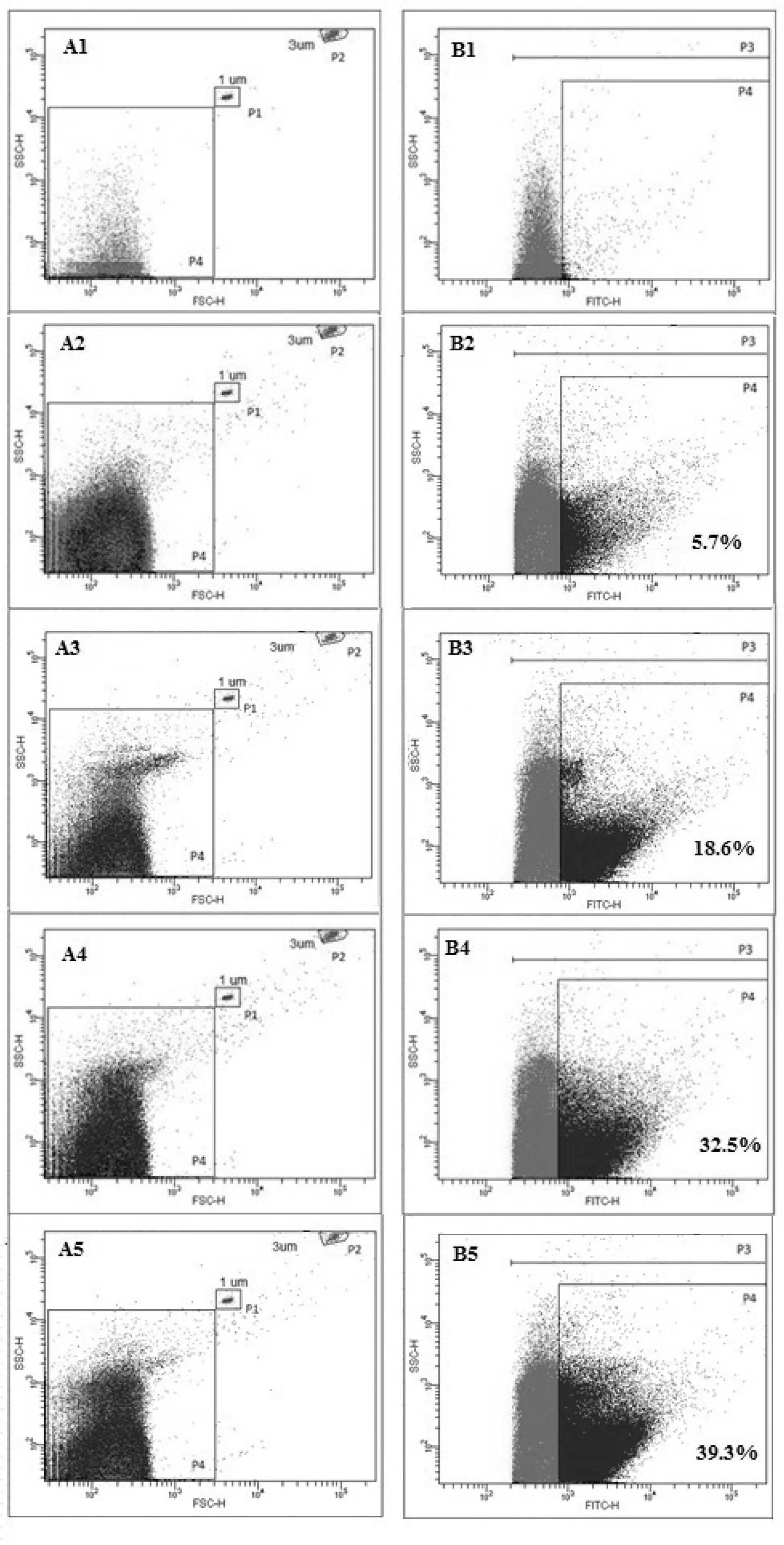
Analysis and counting of MPs of different cellular origins by flow cytometry on a FACS Canto The noise threshold was set up in the FITC fluorescence channel (200 a.u) (B1-B5). A1, B1 – filtered HBS buffer without MPs (negative control); A2, B2 – MPs from erythrocytes; A3, B3 – MPs from platelets; A4, B4 – MPs from THP-1 cells; A5, B5 – MPs from ECs. Annexin V-FITC was added to all probes. All events above the FITC fluorescence threshold (200 a.u) (gate P3 in B1-B5) were counted in the SSC/FSC window (A1-A5) in the size gate < 1 µm (gate P4). After subtracting the events in the negative control, all events in this gate were considered MPs. Gate P1 (A1-A5) – size calibration beads (1 µm), gate P2 (A1-A5) – counting beads (3 µm). Gate P4 (B1-B5) annexin V-positive events above the threshold set up in the negative control in the FITC fluorescence channel. Percentages of annexin V-positive events are presented for each type of MPs. Analysis of MPs from monocytes is not shown since they have approximately the same distribution pattern as MPs from monocytic THP-1 cells (approximately 30% of annexin V-positive events).

### Preparation of phospholipid vesicles

Artificial phospholipid vesicles composed of 80% phosphatidylcholine (PC) and 20% phosphatidylserine (PS) or 15% PS and 85% PS or 10% PS and 90% PC were prepared according to the protocol recommended by Avanti Polar Lipids with minor changes. Phospholipids dissolved in chloroform were transferred into a round-bottomed flask, dried for 30 minutes under a nitrogen current to eliminate chloroform, and hydrated in 20 mM HEPES, 140 mM NaCl (pH 7.5) buffer for 30 minutes at 55°C on a shaker. The resulting solution was treated with three freeze-thaw cycles, heated to 55°C, and forced through the extruder membrane. The pore diameter was 100 nm.

### Plasma preparation

Blood from healthy donors was collected into Greiner Bio-One Vacuette or Sarstedt Monovette citrate tubes. The first tube collected after the venipuncture was discarded. Platelet-free plasma was obtained by two-stage centrifugation: 15 min at 1600 *g* and 5 min at 10,000 *g*. MPs were removed by centrifugation for 30 min at 16,000 *g*. Experiments were carried out on unfrozen plasma of individual donors.

### Thrombodynamics test

The procoagulant activity of MPs was studied using the Thrombodynamics test. The assay is described in [33–35]. A total of 107 µl of MP-poor plasma was supplemented with 13 µl of MPs in different concentrations or buffer A. The assay was performed using a Thrombodynamics Analyzer and Thrombodynamics kit (LLC HemaCore, Moscow, Russia): 120 µl of plasma supplemented with MPs was transferred to an Eppendorf tube containing corn trypsin inhibitor (CTI), incubated for 3 min at 37°С and then transferred to an Eppendorf tube containing Ca salt. Recalcified plasma was placed into the chamber. An insert with immobilized TF was immersed into plasma. Clot growth began from the TF-covered surface. It was monitored by light scattering using a digital camera for 60 min. Clot images were used to determine the clot size, measured as the distance from the edge of the activator to the point where the light scattering intensity was half of the maximal light scattering of the clot at the activator. Clot growth rates were determined as the slope of the clot size dependence on time within the interval from 2 to 6 min after the beginning of clot formation (initial rate, Vi) and 15 to 25 min (stationary rate, Vst).

The appearance of clotting centres within a distant from the activator (spontaneous clots) was characterized by Tsp, the time when the plasma volume excluding clot growing from the activator clotted to 5% (Fig. 2).

**Fig. 2.**
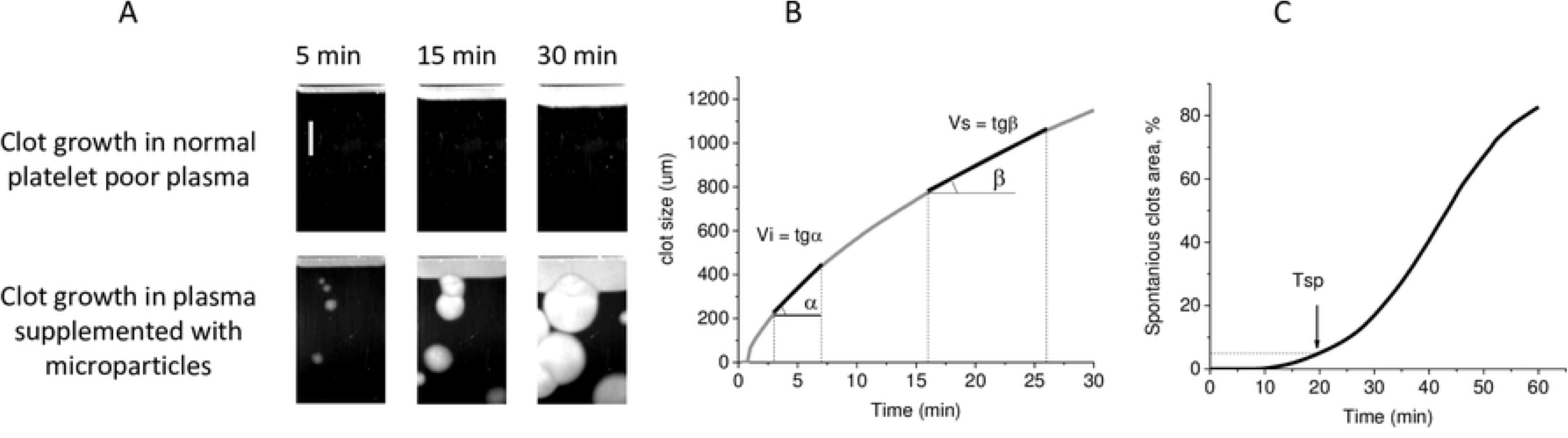
Design of the Thrombodynamics assay. A. Typical images of growing fibrin clot in normal platelet-free plasma and in plasma supplemented with microparticles. Coagulation is activated by immobilized TF (on the top), and the fibrin clot grows into the bulk of the plasma. When plasma is supplemented with microparticles, spontaneous clots in the bulk of the plasma appear. The scale bar is 2 mm. B. Clot size dependence on time, definition of initial clot growth rate, Vi, and stationary clot growth rate, Vs. C. Dependence on time of the percentage of spontaneous clot area of the whole chamber area excluding walls and clot grown from activator (Tsp definition).

To calculate a number of spontaneous clots dependence on time, the photos were converted into binary format. Black colour corresponded to regions with a liquid state of plasma, and white colour corresponded to regions with fibrin clots. The transition occurred when the light scattering intensity at a point became higher than half of the maximal light scattering intensity of a clot growing from the activator. The time point when a spontaneous clot area exceeded an arbitrarily chosen value of 9 µm^2^ was defined as the time of spontaneous clot appearance.

To characterize the growth of individual clots, we introduced the following parameters: lag- time (t lag), maximal rate of intensity increase (VImax), and the rate of coagulation front propagation from a spontaneous clot centre (Vsc). Spontaneous clots induced by low concentrations of MMPs and EMPs terminated growing early, and their light scattering intensity often did not reach half of the maximal light scattering intensity of a clot growing from activator. Therefore, we defined the lag-time as the time when the light scattering intensity in the centre of a spontaneous clot reached 2000 arbitrary units. VImax was defined as the maximum of the time derivative of light scattering intensity in a spontaneous clot centre. The size of the spontaneous clot was defined as the distance between the spontaneous clot centre and the coordinate, where the light scattering intensity from the clot was half the intensity at the centre of the clot. The clot size dependence on time was calculated until fusion of the clot with a neighbouring clot or until the end of the test. Vsc was calculated as the linear approximation of the last 10 min of clot size dependence on time.

### Preparation of fVIIai

The fVIIa inactivation method was carried out as described in [36] with modifications. FVIIa was incubated with PPACK for 60 min at 4°C at a 1:2 molar ratio. The obtained fVIIai was separated from PPACK by dialysis against Tris buffer. The final concentration of fVIIai was measured on a spectrophotometer by absorption at 280 nm. A concentration of 50 nM fVIIai completely suppressed clotting caused by 0.25 pM TF in a 2-hour experiment. Complete inactivation of fVIIa was verified with a chromogenic assay. The FVIIai sample and calibration fVIIa samples 10 ÷ 1.25 pM dissolved in Ca solution were incubated for 5 min with TF, mixed with fX and incubated for 15 min. The reaction was stopped by EDTA addition, and the amount of fXa gained was evaluated by the cleavage rate of the substrate S 2765. PPACK removal was checked by the influence of fVIIai solution on fluorogenic substrate cleavage by thrombin.

### Determining the activation pathway from MPs

To determine the activation pathway from MPs, the time of spontaneous clots appearance (Tsp) in plasma supplemented with MPs of different types was measured without any inhibitors, with the addition of 200 µg/ml CTI, 100 nM VIIai (TF pathway inhibitor) or both inhibitors.

## Results

### MPs of different origins in coagulation activation

The ability of MPs to activate coagulation can be observed in the Thrombodynamics test by the appearance of clotting centres at a distance from the activating surface. These clots are called spontaneous because their appearance is caused by material in the plasma and not by activation with substances added in the test.

MPs of different origins were titrated in MP-depleted plasma in the Thrombodynamics test. MP concentrations varied from 0 to the value at which clotting centres in the plasma volume appeared within 60 min. Spontaneous clotting parameters induced by MPs of different origins turned out to have both quantitative and qualitative differences.

The minimal MPs concentrations causing spontaneous clotting could serve as the measure of MPs activity in coagulation activation. The data for MPs of different origins are represented in Table 1. The activity of MMPs from cells of normal donors was the highest. The activity of MPs from monocyte culture THP cells was 4-fold lower. EMPs had activity 25-fold less than that of MMPs. PMPs and ErMPs were 100-fold less. Although the deviation between the activity of MPs samples obtained from cells of different donors reached from 48% for PMPs to 98% for ErMPs, all the differences between MPs activity were significant (р<0.05), except for the difference between PMPs and ErMPs.

**Table. 1.**
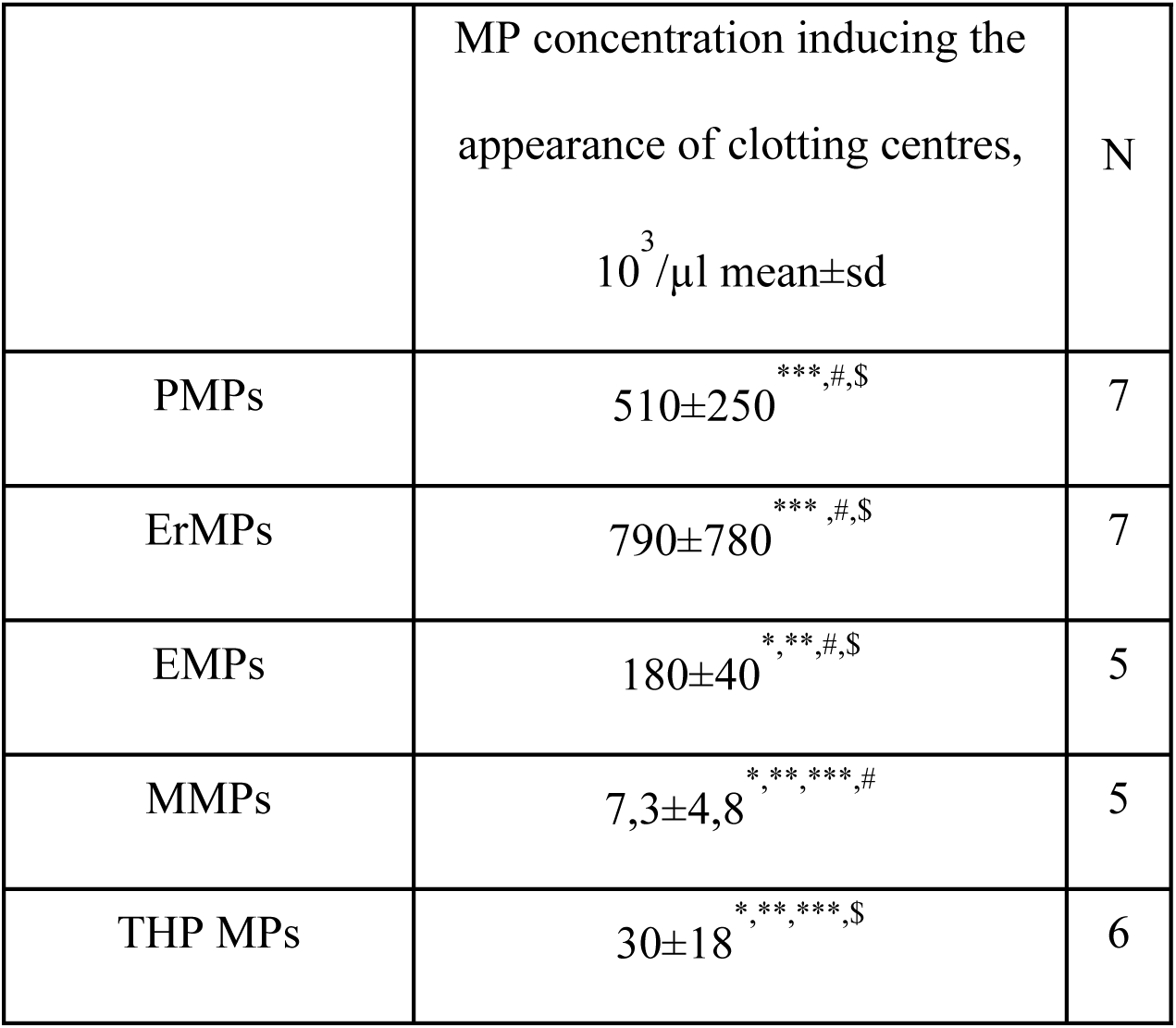
The minimal MPs concentrations inducing the appearance of clotting centres. For analysis of significant differences, the Mann-Whitney test was used. * p<0.05 – significant difference with PMPs, ** p<0.05 – significant difference with ErMPs, *** p<0.05 – significant difference with EMPs, # p<0.05 – significant difference with THP MPs, $ p<0.05 – significant difference with MMPs. N – the number of MPs samples tested.

Even more evident than the minimal MPs concentration causing spontaneous clots, the difference in the MPs activity was observed in the ratio of the number of coagulation centres formed within 1 hour to the total number of MPs in the chamber. This parameter depended on MPs concentrations; however, the differences between MPs of different origins were so great that even the range from the 25th to the 75th percentile from the complete range of this ratio change at different concentrations of MMPs, EMPs and PMPs did not intersect (Table 2). For the most active MMPs, less than 0.7 per thousand MMPs caused a visible centre within 1 hour. Less than 1/10^5^ EMPs and 1/10^6^ PMPs or ErMPs induced a separate clot.

**Table. 2.**
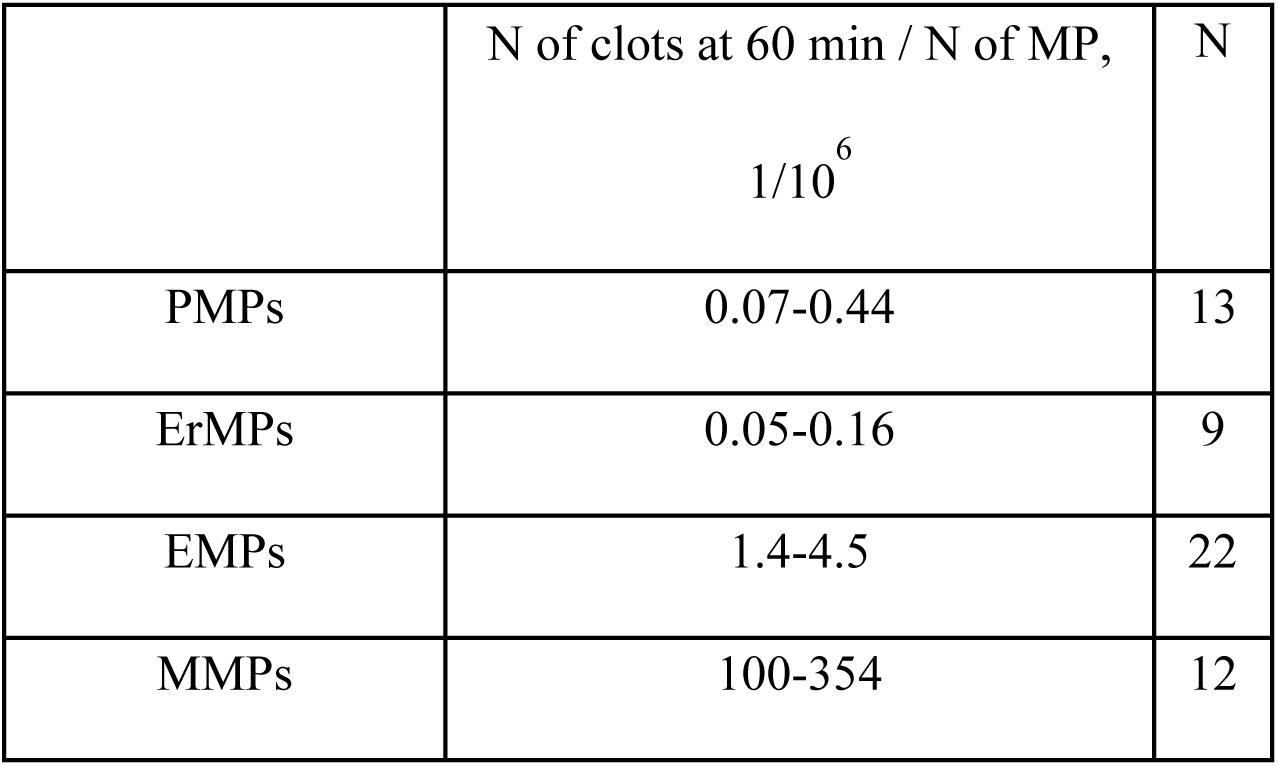
The ratio of spontaneous clots number to the number of MPs in the chamber volume. The number of MPs in the chamber volume was estimated based on MPs concentrations and considering the plasma volume where spontaneous clots were counted to be equal to 70 µl. N – the number of tests included in the calculation. The ranges of values in the table are from the 25th to 75th percentile.

| | N of clots at 60 min / N of MP,<br>$1/10^6$ | N |
| --- | --- | --- |
| PMPs | 0.07-0.44 | 13 |
| ErMPs | 0.05-0.16 | 9 |
| EMPs | 1.4-4.5 | 22 |
| MMPs | 100-354 | 12 |

One of the reasons for the activity difference could be TF on the MPs surfaces. We determined the activation pathway from MPs by the changes in the time of spontaneous clots appearance Tsp in recalcified plasma containing MPs of different origins when inhibitors of contact activation (CTI) or the TF pathway were added. The test showed that the most active MMPs and EMPs bear TF on their surfaces (Fig. 3). In addition to TF, contact activation made a significant contribution to EMPs activity. Less active ErMP and PMP activate coagulation through the contact pathway only (Fig. 3).

**Fig. 3.**
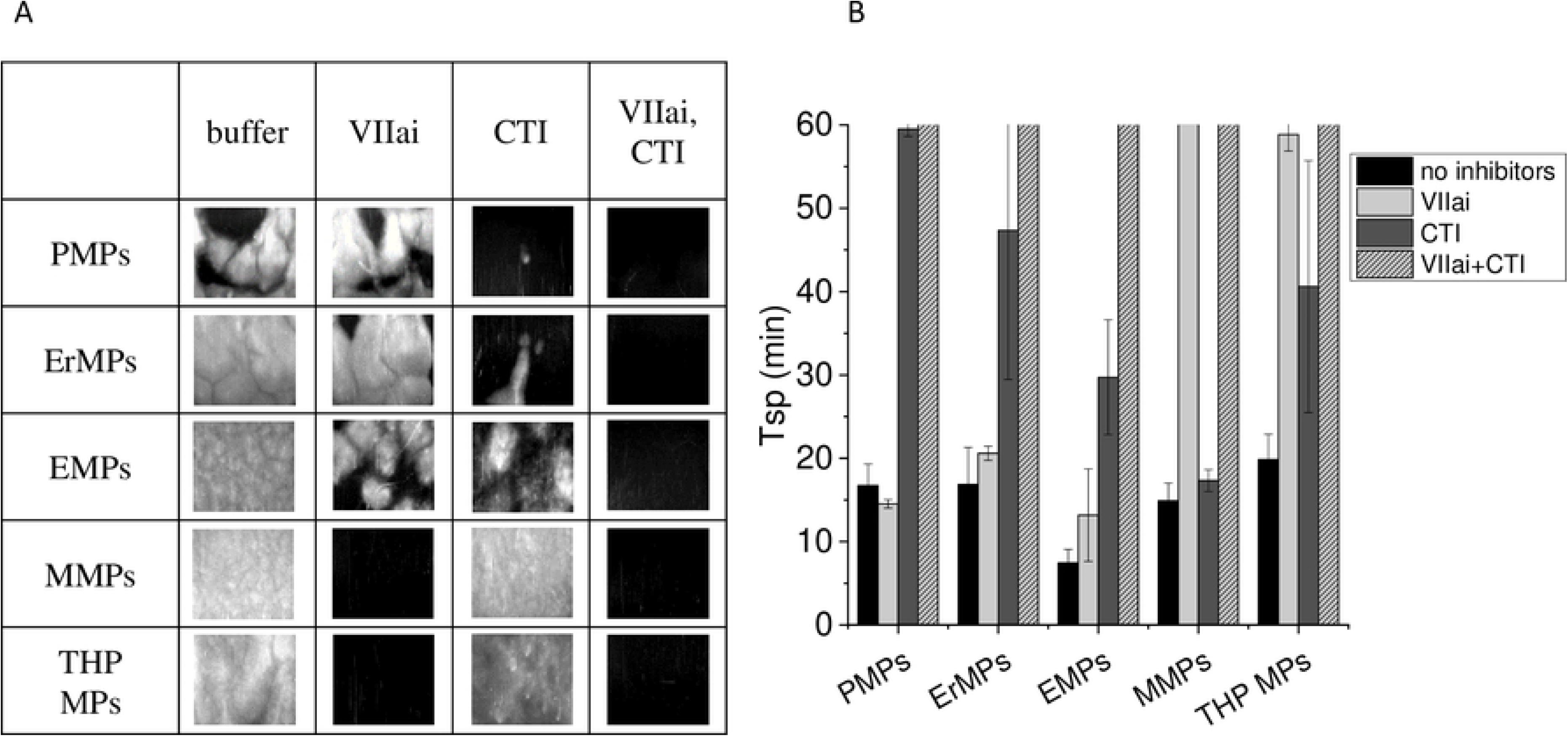
Activation pathway from MPs. Photos at 60 min of plasma supplemented with MPs of different origins in the Thrombodynamics test without any inhibitors, with 100 nM VIIai (TF pathway inhibitor), with 200 µg/ml CTI (contact pathway inhibitor) or both inhibitors. MPs were supplemented at arbitrary concentrations, which induced the appearance of clotting centres within 10-20 min in samples without inhibitors. That was optimal for checking inhibitor effects. B. Mean ± sd of Tsp in plasma supplemented with MPs without inhibitors and with one or both inhibitors. Three repeats were carried out for PMPs and THP MPs and two repeats for ErMPs, EMPs and MMPs. PMPs – platelet microparticles, ErMPs – erythrocyte microparticles, EMPs – endothelial microparticles, THP МPs – microparticles from monocyte culture, MMPs – monocyte microparticles.

Since the minimal concentrations leading to the appearance of spontaneous clots for MPs of different origins differ up to a hundred times, the typical patterns of spontaneous clots formation are presented in individual concentrations for each type of MPs (Fig. 4, S1 Movie). Typical coagulation patterns had qualitative differences in the number, appearance and growth rate of spontaneous clots.

**Fig. 4.**
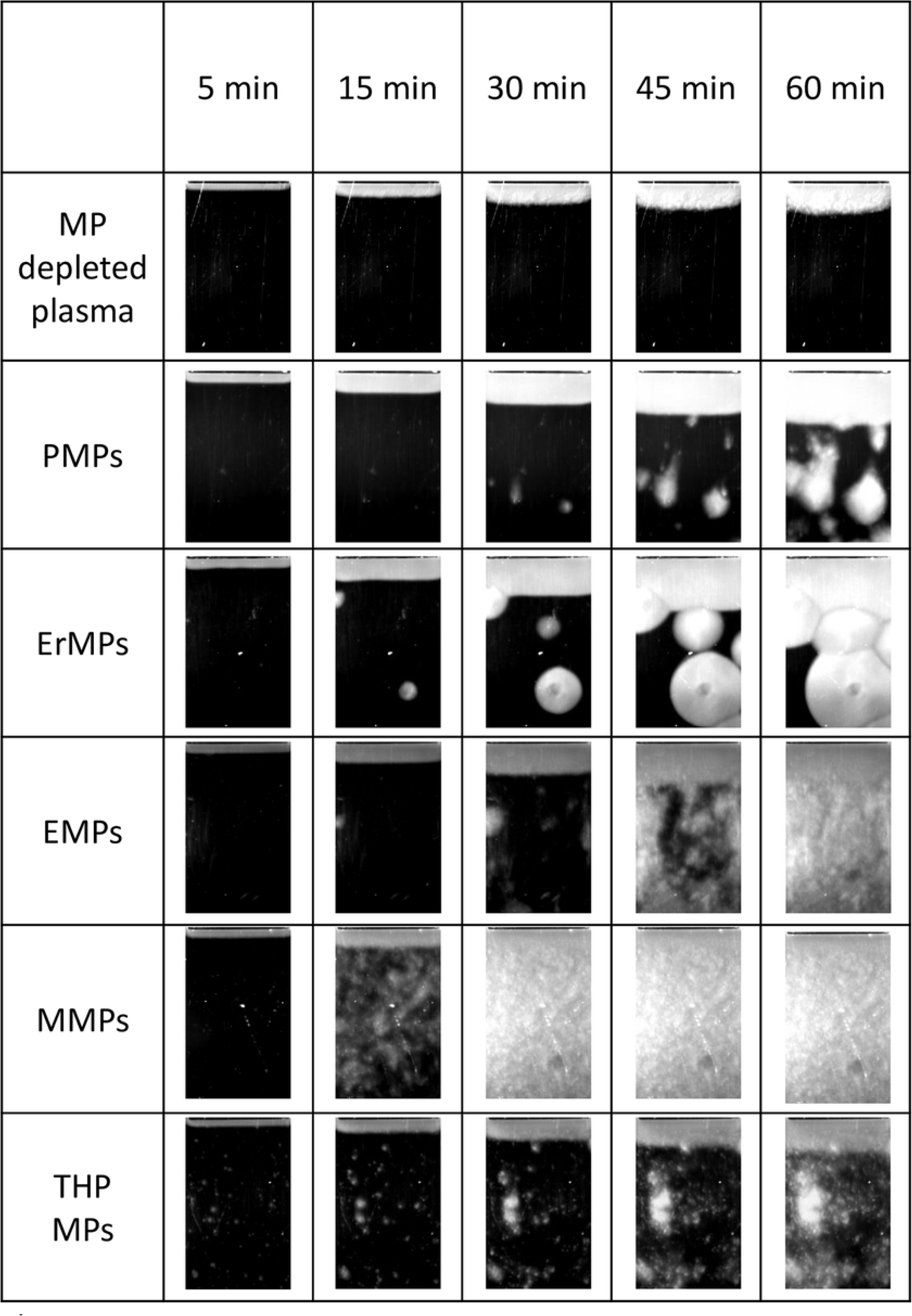
Photos of growing clot and typical patterns of spontaneous clotting induced by MPs of different origins. As MPs of different origins induce spontaneous clotting at 100-fold different concentrations, the photos represented at arbitrary concentrations at which the patterns of clotting centre appearance were well distinguished within 60 min. The MPs concentrations in assays in photos were as follows: PMPs, 627·10^3^ 1/µl; ErMPs, 500·10^3^ 1/µl; EMPs, 480·10^3^ 1/µl; THP MPs, 132·10^3^ 1/µl; and MMPs, 132·10^3^ 1/µl.

The time dependences of the number of clotting centres induced by MPs of different origins are represented in Fig. 5. The increment of the clot number was normalized to the fraction of free frame area so that the decrease in the rate of clotting centre appearance as a result of volume occupation by clots that appeared earlier would not distort the dependence. The scale of the ordinate axis in Fig. 5 is different because the number of spontaneous clots induced by MPs of different origins within 1 hour differed by more than 100-fold. According to available data for PMPs and ErMP, the form of the clot number dependence on time is difficult to determine. For MMPs and EMPs, this dependence was exponential. It is natural to assume that the time of appearance of a spontaneous clot is determined by the local concentration of the activator in its centre. If an individual MP triggered a clot, the time of appearance of the first five centres (t_N=5_) should not depend on the MPs concentration. For ErMP, the assumption was confirmed (Fig. S1 B). For PMPs and EMPs, t_N=5_ tended to decrease with increasing MPs concentration, but the data were not sufficient for a definite answer (Fig. S1 A, C). For MMPs, t_N=5_ decreased inversely with MMPs concentration (Fig. S1 D). This indicates an increase in the probability of clot formation with a decrease in the distance between MMPs. The assumption about the interaction between MPs is also supported by the fact that for all types of MPs in experiments where more than 10 centres were formed within 1 hour, the increase in the number of centres was accelerated with time.

**Fig. 5.**
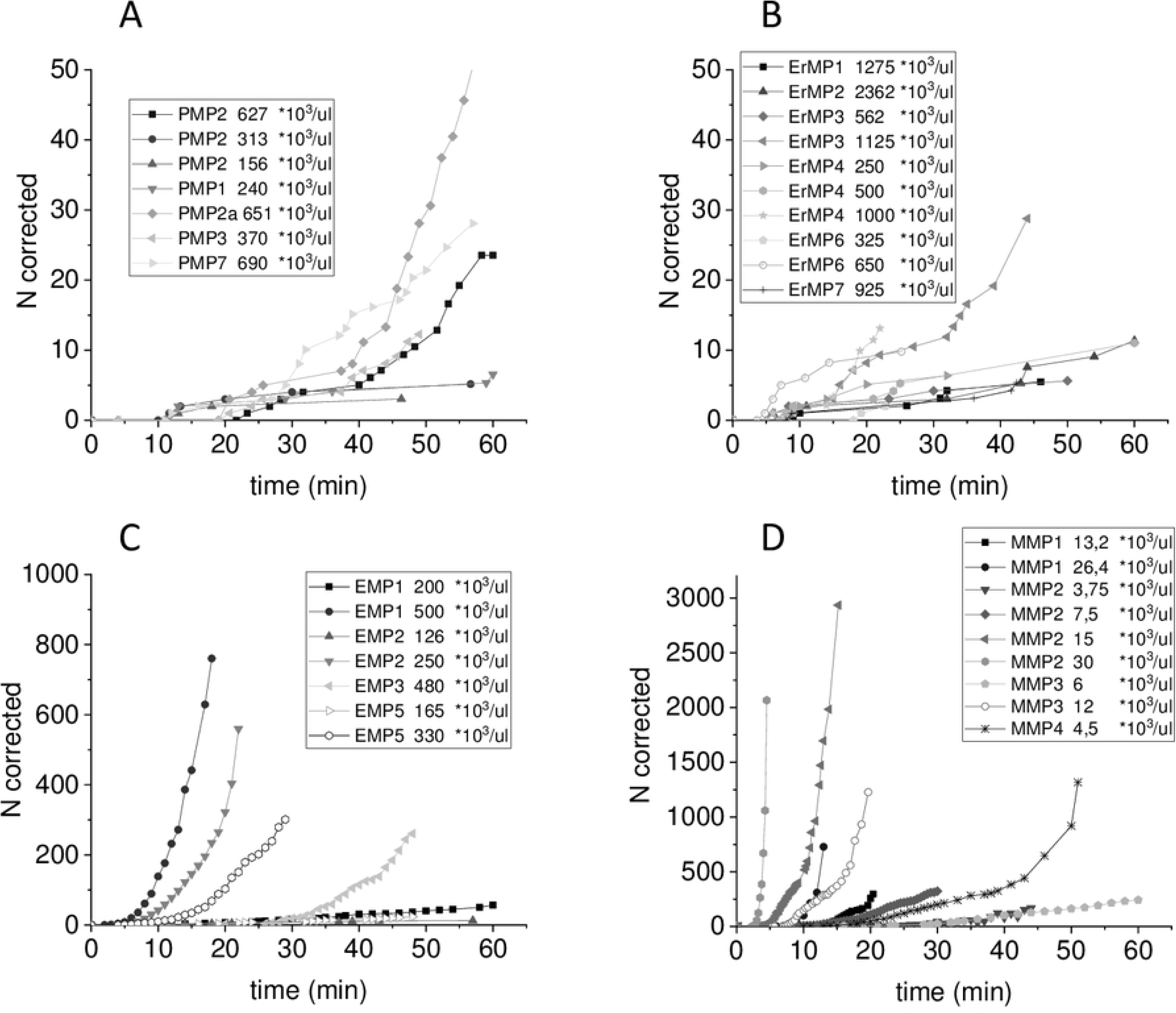
Time dependence of the number of clotting centres induced by MPs of different origins. The number of clotting centres was recalculated to represent what it would have been if the plasma volume had not been decreased by clots that appeared earlier (N corrected). Clotting was induced in normal MP-depleted plasma by supplementation with (A) platelet MPs, (B) erythrocyte MPs, (C) endothelial MPs, and (D) monocyte MPs. Different curves correspond to different MPs samples and different concentrations as described in the legends.

The dependences of the number of clotting centres formed within 60 min on concentrations are represented in Fig. 6. ErMPs in minimal concentrations that induce spontaneous clotting led to the appearance of few spontaneous clots growing at a rate near the rate of clot growth from the surface with immobilized TF (activator). On average, this rate was 48±14 µm/min. The number of spontaneous clots increased along with the ErMP concentration, but because of their fast growth, they rapidly occupied all the free chamber volume, and in our experiments, the maximal number of spontaneous clots formed within 1 hour did not exceed 17 (Fig. 6 B). Clots induced by PMPs grew at a mean rate of 14±8 µm/min. As a result, more clots could appear at high PMPs concentrations before full coagulation (Fig. 6 A). MMPs with concentration increase did not cause coagulation at first then approximately ten clots appeared. The growth rate of these clots almost stopped within the first 10 – 15 min, and the whole plasma volume did not clot. With a further slight increase in concentration, there was a sharp switch to the formation of several hundred clots (Fig. 6 D) growing at a relatively low but significantly nonzero rate of 9±4 μm/min on average. In this case, plasma coagulated completely and not only because of individual clots propagation in space but also because of coagulation in the whole volume. EMPs caused spontaneous coagulation patterns qualitatively similar to MMPs, but the dependence of the number of spontaneous clots on concentration was considerably smoother (Fig. 6 C), and at high EMPs concentrations, much higher clots growth rates were achieved.

**Fig. 6.**
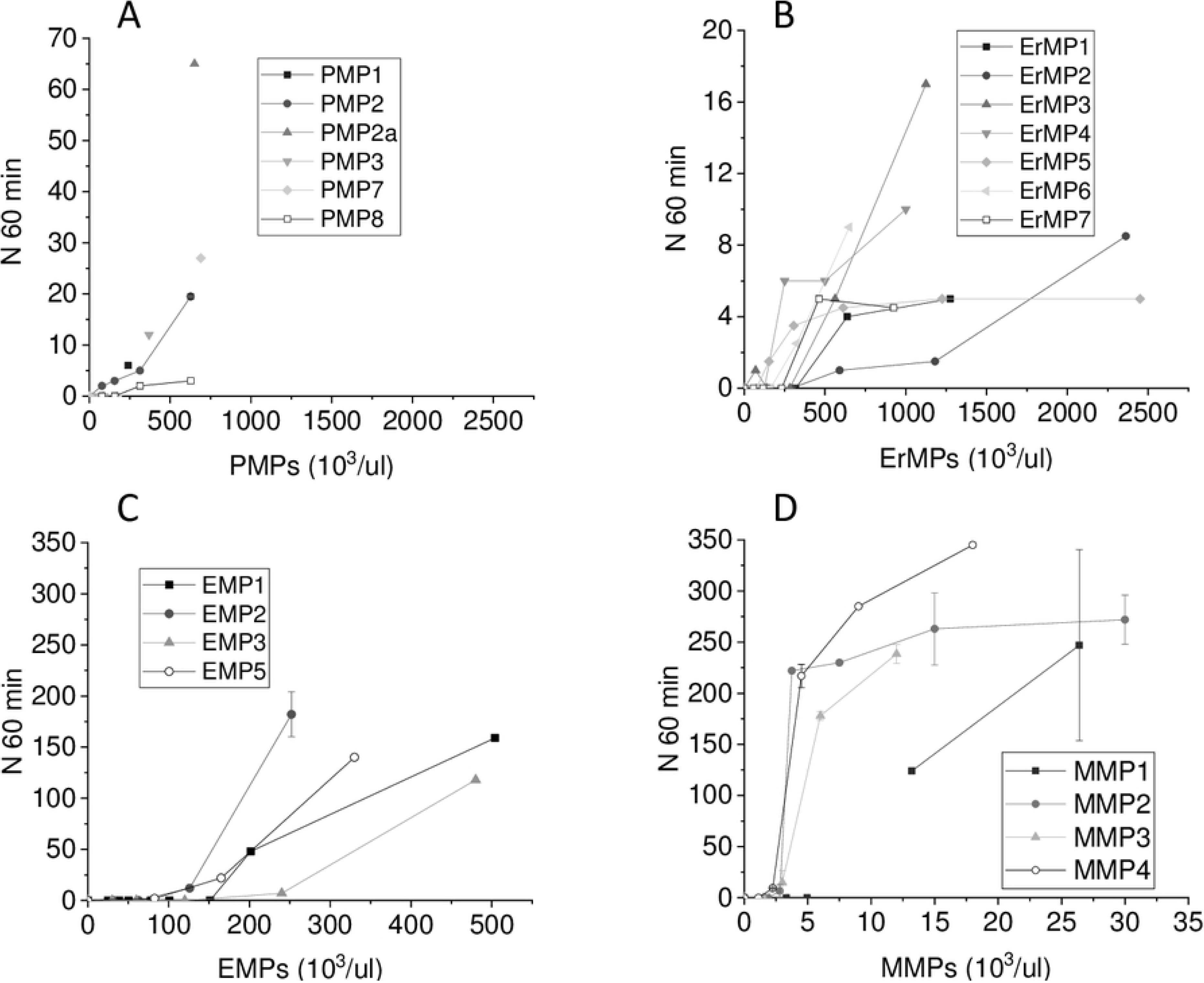
The dependence of the number of clotting centres formed within 60 min on concentration. Clotting was induced in normal MP-depleted plasma by supplementation with (A) platelet MPs, (B) erythrocyte MPs, (C) endothelial MPs, and (D) monocyte MPs.

As described above, the distribution of fibrin clots in space in this method is reflected by the light scattering intensity profile. Analysis of the time dependences of light scattering intensity profiles allows the introduction of quantitative characteristics of spontaneous clots. The corresponding dependences when MPs of different origins were supplemented to plasma are represented in Fig. 7. PMPs and ErMPs induced clotting in separate centres only. The light scattering intensity in the centres of clots increased along with fibrin polymerization. The area with high light scattering intensity grew as the clot propagated in space (Fig. 7 A, D). The rate of light scattering increase in the clot centre was characterized by the maximal rate VI max and the coagulation front propagation from centres of spontaneous clots with the rate Vsc. VI max and Vsc were calculated as described in the materials and methods. If all the fibrinogen cleaved to fibrin within the test time, the light scattering intensity in the clot centre reached a plateau and only further propagation in space continued. The maximal light scattering intensity in a clot centre reached by 60 min was indicated as Imax. With increasing MMPs and EMPs concentrations, qualitative changes in the parameters of spontaneous clots growth were observed. Therefore, the time dependences of the light scattering intensity profiles of spontaneous clots induced by MMPs and EMPs are given for two concentrations, conditionally “low” and “high”. In plasma containing MMPs and EMPs in “high” concentrations, coagulation began in the whole volume at some point in time. That was expressed as the background increase in light scattering intensity (Fig. 7 С, F). At conditionally “low” MMPs concentrations, distinct clotting centres appeared, the growth of which rapidly stopped (Fig. 7 E). EMPs were especially heterogeneous: in the same sample at conditionally “low” concentration, coagulation propagated from some centres and practically stopped from others (Fig. 7 B).

**Fig. 7.**
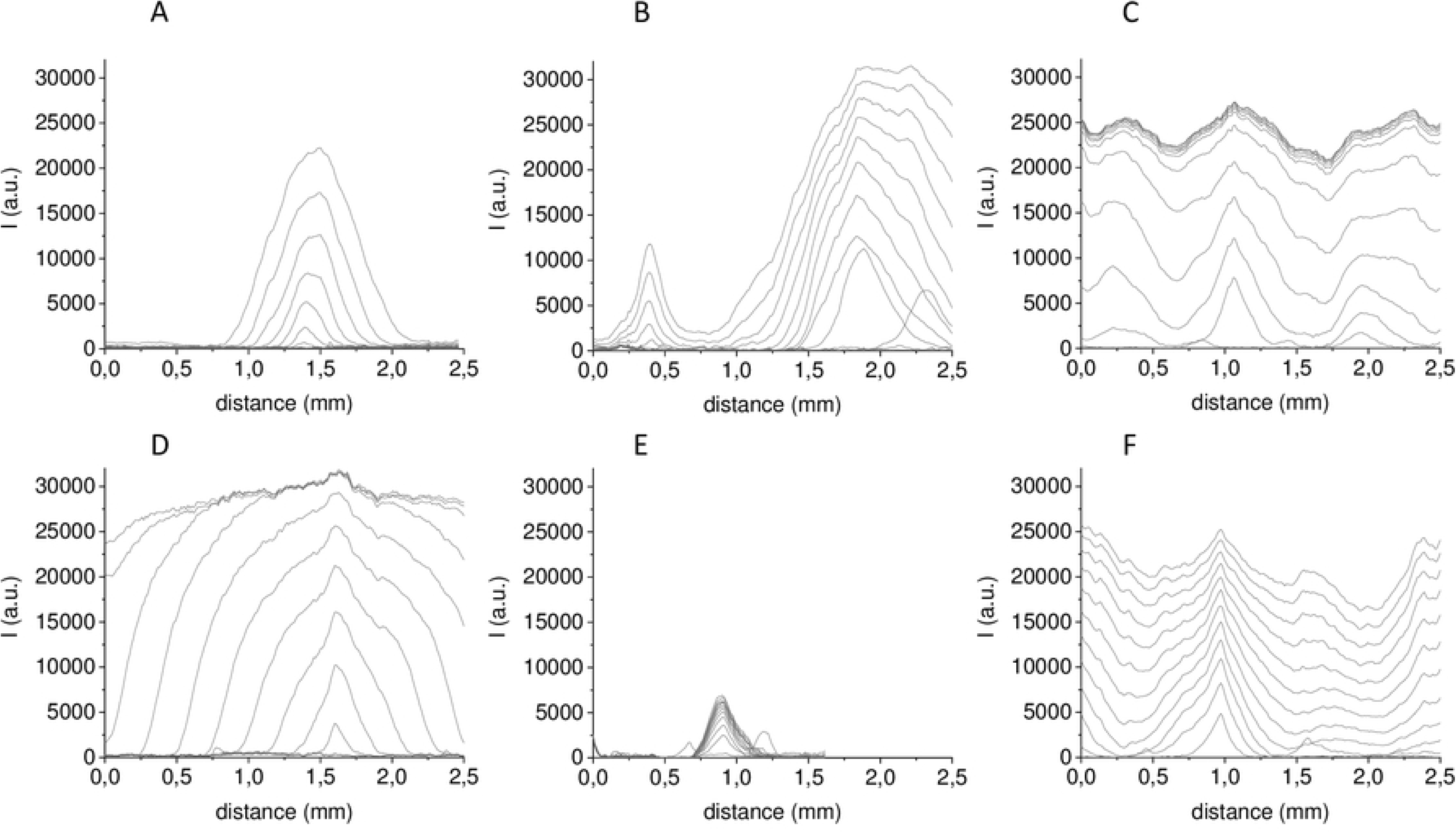
The light scattering intensity profiles of spontaneous clots induced by MPs of different origins. Clotting was induced in normal MP-depleted plasma by supplementation of (A) platelet MPs, (B) endothelial MPs in conditionally “low” concentrations, (C) endothelial MPs in conditionally “high” concentrations, (D) erythrocyte MPs, (E) monocyte MPs in conditionally “low” concentrations, and (F) monocyte MPs in conditionally “high” concentrations. The time interval between profiles is 5 min.

VI max is by some approximation proportional to the rate of fibrin formation and consequently to the thrombin concentration (at the time interval until complete polymerization). Therefore, VI max indirectly indicates activation strength. VI max in the centres of spontaneous clots induced by any type of MPs was considerably less than VI max at the surface with immobilized TF (activator) (Fig. 8). At the same time, Imax in the centre of spontaneous clots could exceed Imax at activator by 1.4 – 1.7- fold (Fig. 8, S2). The increase in maximal light scattering intensity of a clot as a result of a diameter increase of fibrin fibrils when the thrombin concentration is decreased was theoretically predicted and experimentally shown in the article [37]. The VI max deviation in one test consisted of 33% for PMPs and 60% for MMPs. The VI max changes with concentration were less than the deviation within a test for all types of MPs except for EMPs (Fig. S3), which gives justification for a rough approximation to compare the average VI max for all clots formed at different concentrations. The comparison is shown in Fig. 9 A. The mean VI max of ErMPs is significantly higher than the mean VI max of the other types of MPs, and for MMPs, it is significantly lower. PMPs and EMPs did not differ significantly in this parameter. However, the VI max in the centre of spontaneous clots induced by any type of MPs was not higher than the 0.1 – 0.15 VI max at activator (Fig. 9 B). Consequently, the thrombin concentration in the centres of spontaneous clots was 7–10-fold lower than that at activator.

**Fig. 8.**
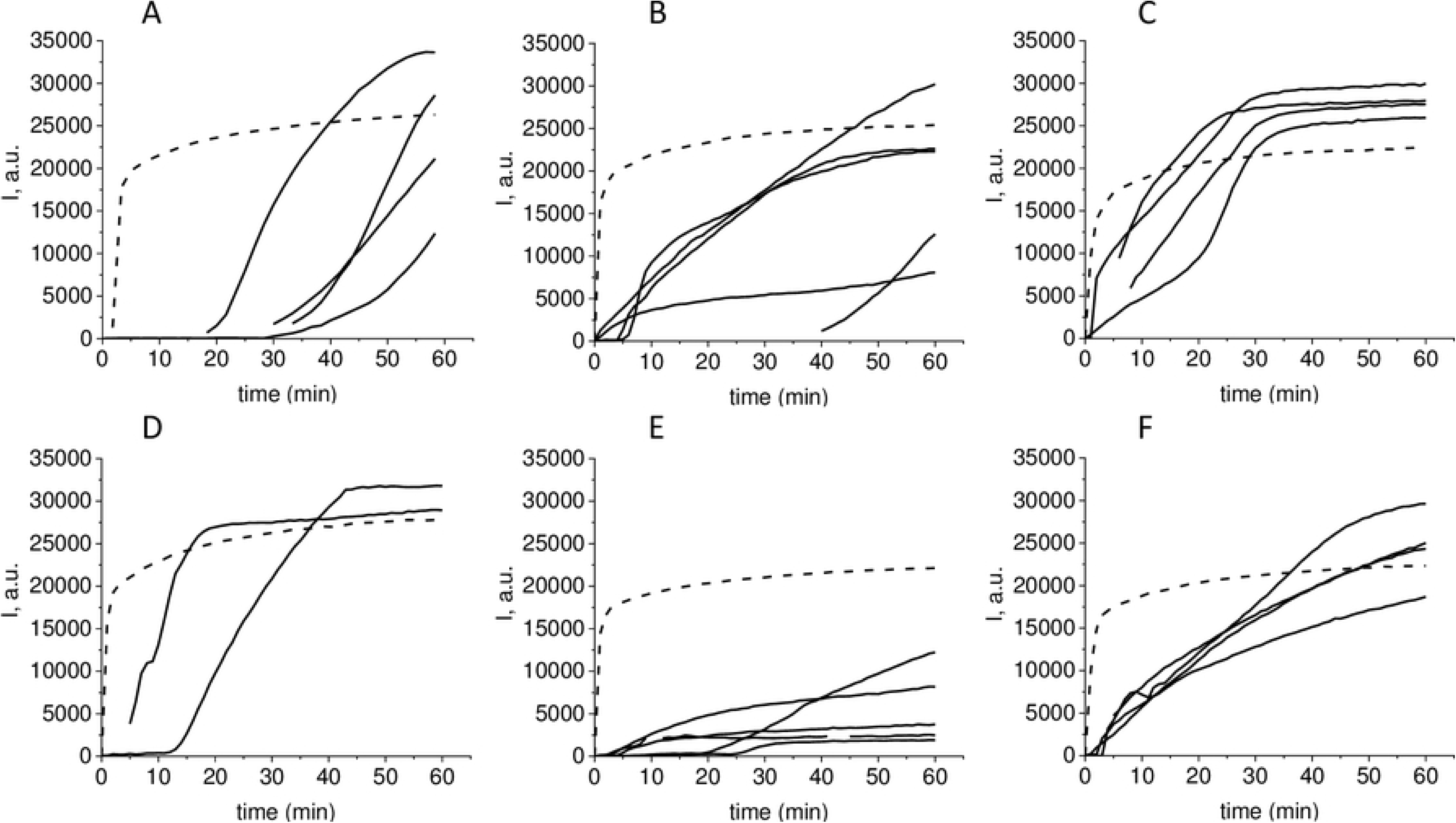
Time dependence of light scattering intensity in the centre of spontaneous clots and clots growing from activator. Clotting was induced in normal MP-depleted plasma by supplementation of (A) platelet MPs, (B) endothelial MPs in conditionally “low” concentrations, (C) endothelial MPs in conditionally “high” concentrations, (D) erythrocyte MPs, (E) monocyte MPs in conditionally “low” concentrations, and (F) monocyte MPs in conditionally “high” concentrations. Time dependence of the light scattering intensity of clots growing from activator are drawn with dashed lines, and those of the light scattering intensity in the centre of spontaneous clots are drawn with solid lines.

**Fig. 9.**
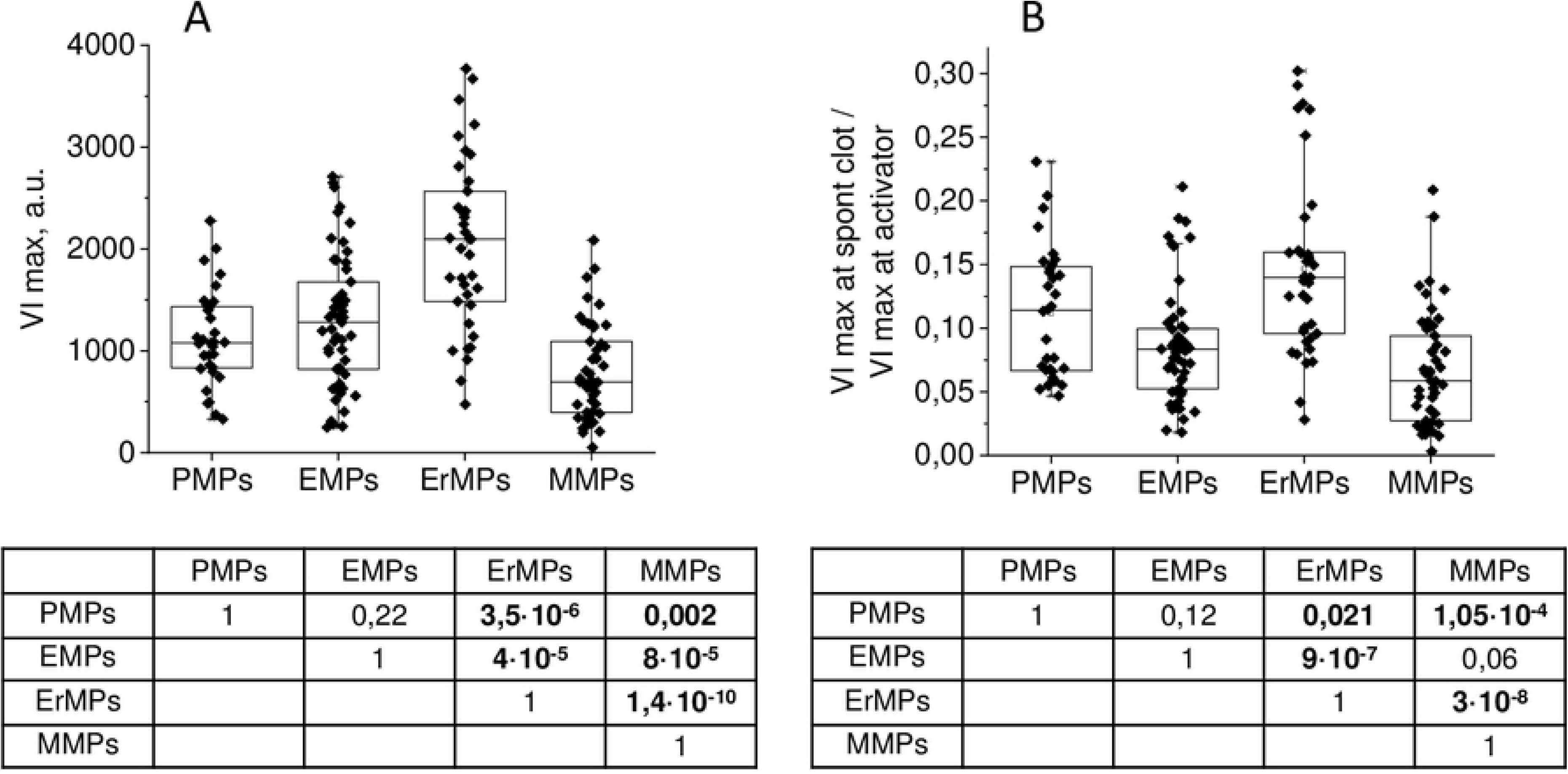
The maximal rate of increase of light scattering intensity in the centre of spontaneous clots and its ratio to the maximal rate of increase of light scattering intensity of clots growing from activator. (A) The mean maximal rate of light scattering intensity increase in the centre of spontaneous clots growth. (B) Ratio of the mean maximal rates of light scattering intensity increase in the centre of spontaneous clots growth to the maximal rate of light scattering intensity increase of clots growing from activator. Dots correspond to different MP samples and different concentrations. Tables under histograms contain significance levels of corresponding parameter differences between MP of different origin according to Mann – Whitney test.

If the increment in light scattering intensity in the centre of a clot depends on thrombin formed in the immediate vicinity of the activator, the clot propagation depends on thrombin formed at a distance from the activator as a result of positive feedbacks work. In this way, the parameters VI max and Vsc are largely determined by different reactions. The first characterizes the activation phase, and the second characterizes the propagation phase. The spontaneous clots size in most cases depended linearly on time (Fig. 10). In some cases, the linear region is preceded by a smooth acceleration from zero rate. When the light scattering intensity background is rising, the growth rate was accelerating (Fig. 10 C), although the consideration of the corresponding region as the growth of a separate clot was not quite correct.

**Fig. 10.**
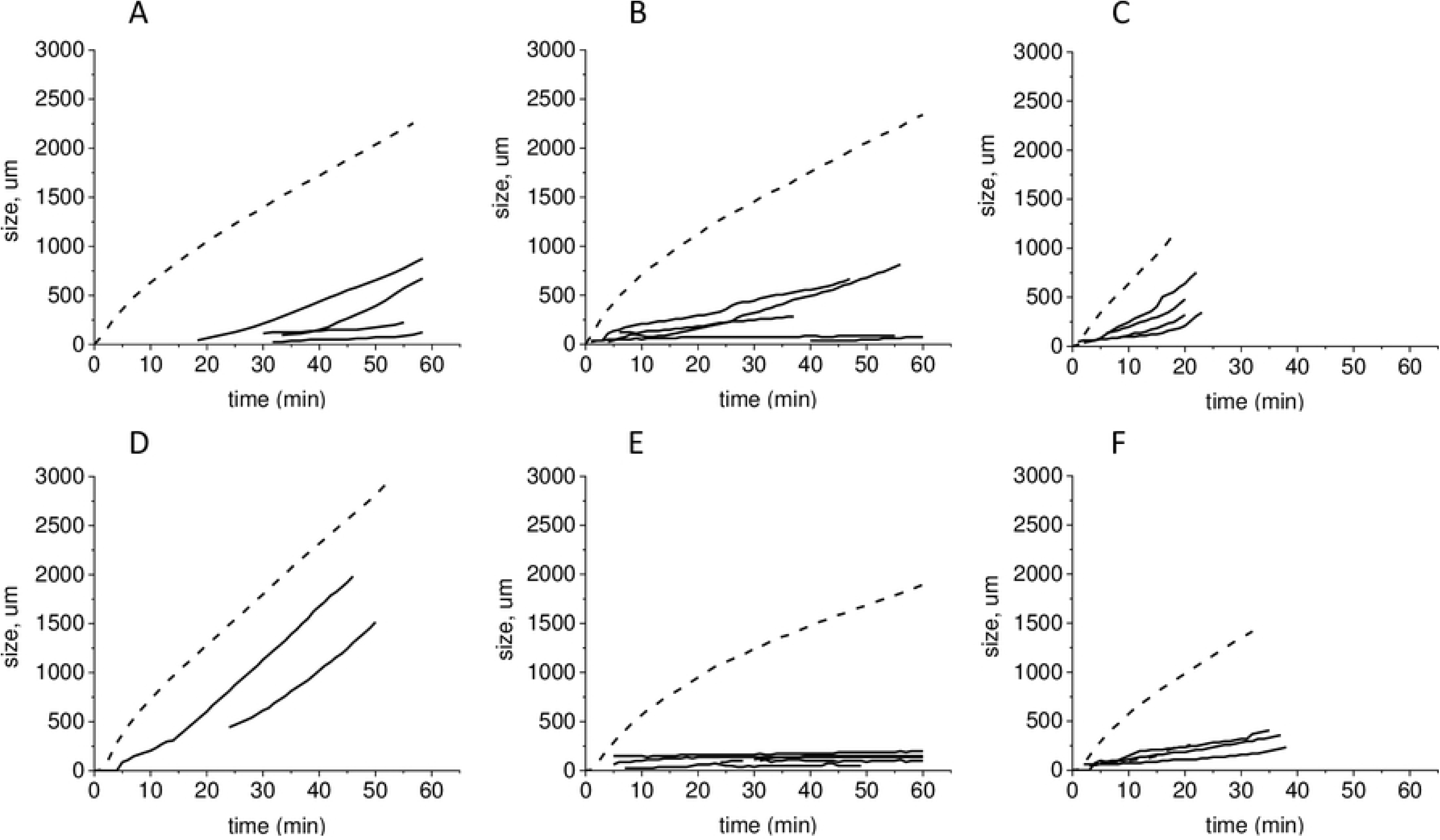
Time dependences of clot growing from activator and spontaneous clots sizes. Clotting was induced in normal MP-depleted plasma by supplementation of (A) platelet MPs, (B) endothelial MPs in conditionally “low” concentrations, (C) endothelial MPs in conditionally “high” concentrations, (D) erythrocyte MPs, (E) monocyte MPs in conditionally “low” concentrations, and (F) monocyte MPs in conditionally “high” concentrations. Time dependences of clots growing from activator sizes are drawn with dashed lines, and those of spontaneous clot sizes are drawn with solid lines.

Growth rates were, on average, increased with concentrations (Fig. S4,) and it would be preferable to compare the growth rates for MPs of different origins at equal concentrations. However, the growth rates of clots induced by TF-bearing MPs cannot be calculated at the concentrations that PMPs and ErMPs begin to initiate clotting, and at concentrations that allow calculating the rates of clots induced by TF-bearing MPs, PMPs and ErMPs do not initiate coagulation. Thus, we represent here a comparison of the parameters of clots induced by MPs of different origins rather than the MPs themselves. In the concentration ranges where clots growth rates could be calculated, changes in the mean rates with the concentrations were less than deviations within one test (the same as for VI max); therefore, we considered the average for different concentrations data (Fig. 11 A). The mean Vsc of clots induced by PMPs and EMPs did not differ significantly. The mean Vsc of clots induced by MMPs was significantly less than that induced by other MPs, and for ErMPs, it was significantly higher than that induced by others. Different phospholipid concentrations due to large differences in MPs concentrations could be one of the reasons for MMPs and ErMPs standing out.

**Fig. 11.**
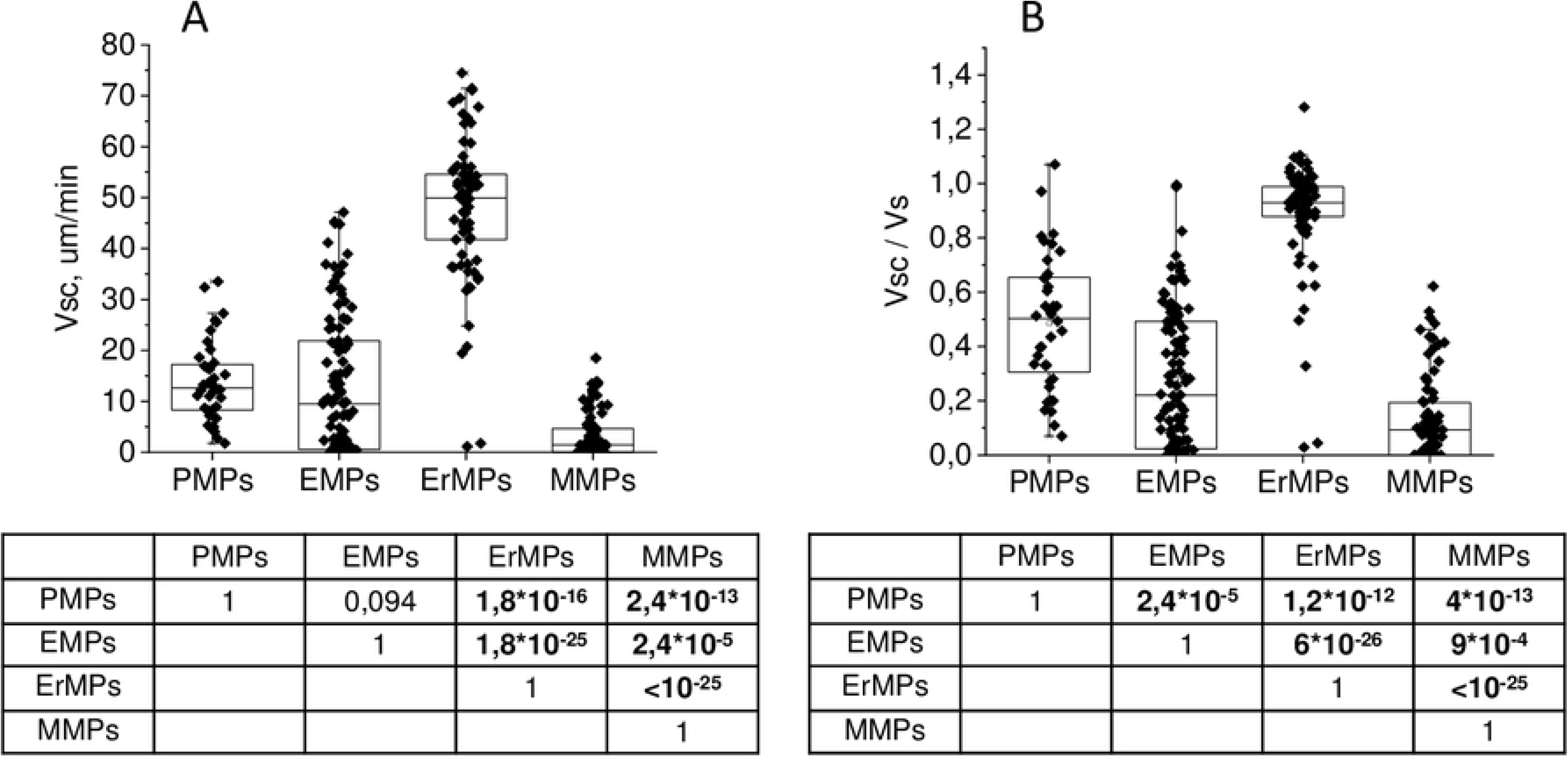
The rate of coagulation front propagation from centres of spontaneous clots induced by MPs of different origin and its ratio to the clot growth rate from activator. (A) Mean rate of coagulation front propagation from centres of spontaneous clots. (B) Mean ratio of the rate of coagulation front propagation from the centres of spontaneous clots to the clot growth rate from the activator. Dots correspond to different MPs samples and different concentrations. Tables under histograms contain the significance levels of corresponding parameter differences between MP of different origins according to the Mann–Whitney test.

It was previously shown that in the presence of a sufficient amount of phospholipid surface in the plasma and activation higher than the threshold, the steady-state clot growth rate over a wide range does not depend on the TF concentration [38], type of TF–bearing cells [39] or the method of activation [40]. One could expect that the growth rates of spontaneous clots and the clot from the activator will be close. However, Vsc deviation within one test consisted on average of 4 µm/min (33%), 8 µm/min (22%), 7 µm/min (76%), and 2.5 µm/min (84%) for PMPs, ErMPs, EMPs, and MMPs, respectively. Only spontaneous clots induced by ErMPs grew at approximately the same rate as the clot from the activator. The medians of the Vsc to Vs ratio were 0.5, 0.93, 0.22 and 0.09 for PMPs, ErMPs, EMPs, and MMPs, respectively (Fig. 11 B). The Spearman correlation coefficients between the growth rates of spontaneous clots and clot from activator were 0.36 (p=0.024) for PMPs, 0.74 (р=3·10^-15^) for ErMPs, 0.78 (р=5·10^-23^) for EMPs, and 0.61 (р=3·10^-10^) for MMPs.

One could assume that any MPs except for ErMPs induce clotting at concentrations that did not supply enough phospholipids to plasma to support coagulation propagation efficiently. ErMPs induce spontaneous clotting at the highest concentration and consequently the highest phospholipid concentration. In the case of lipid deficiency, a dependence of the rate on the activation force of a particular centre can be expected. We did not have the opportunity to measure activation from each centre directly, but indirect data were provided by the rate of increase in light scattering intensity and the lag-time of spontaneous clots appearance. For ErMPs, there was no correlation between VI max and Vsc: r=0.19, p=0.22. For MPs of other types, the correlation of these parameters was significant but weak: 0.43 (p=0.026) for PMPs, 0.58 (p=8·10^-9^) for EMPs, and 0.32 (p=0.004) for MMPs (Fig. 12). The t lag did not show a strong correlation with Vsc also: -0.45 (p=0.03) for PMPs, -0.64 (p=1.2·10^-4^) for ErMPs, -0.35 (p=0.025) for EMPs, and -0.5 (p=0.0015) for MMPs (Fig. 13). Thus, the deviations in Vsc within one test were determined at least not only by the activation strength.

**Fig. 12.**
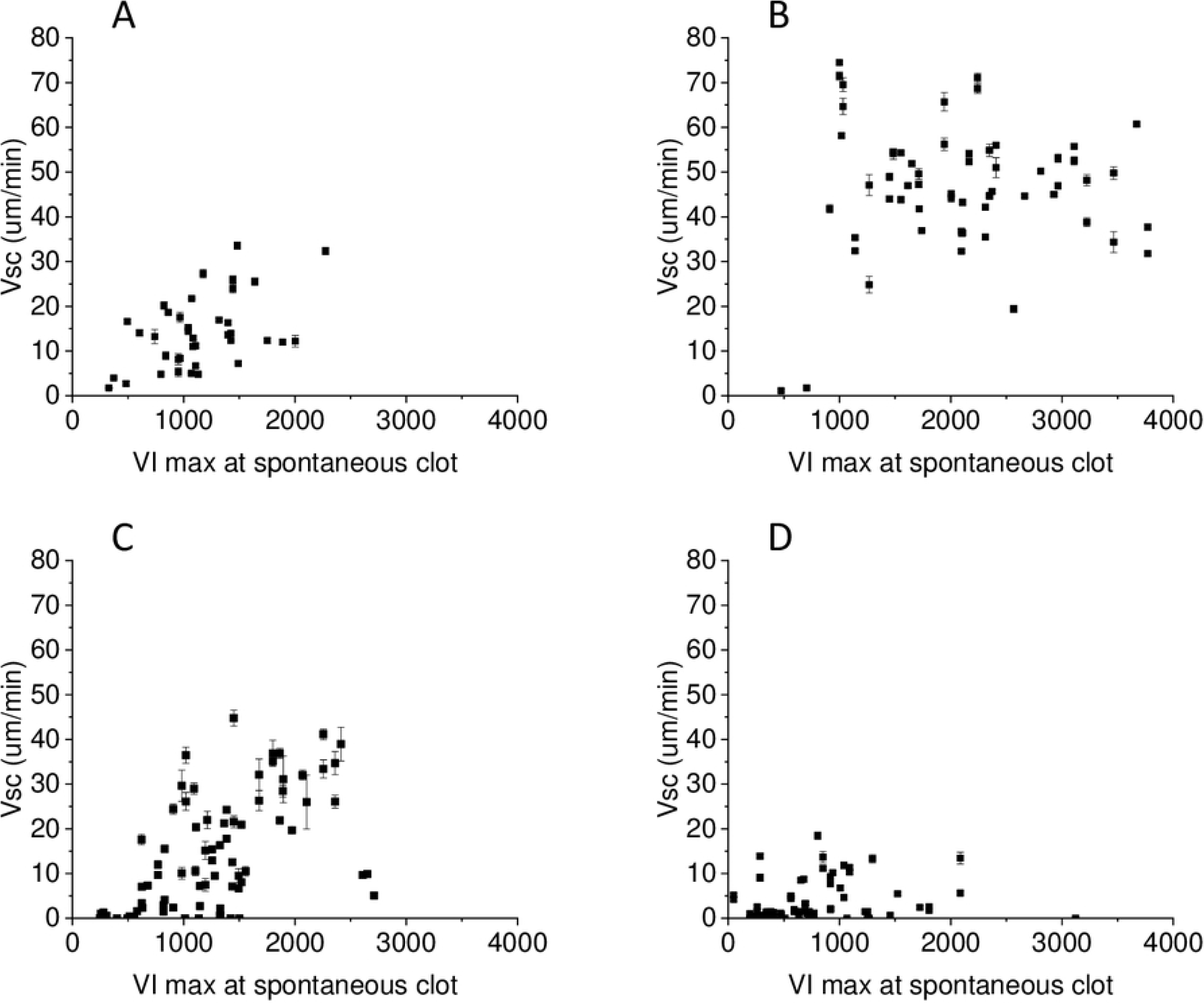
The rate of coagulation front propagation from centres of spontaneous clots and the maximal light scattering intensity in the centre of spontaneous clots growth rate correlation. (A) Data are represented for platelet MPs, (B) erythrocyte MPs, (C) endothelial MPs, and (D) monocyte MPs.

**Fig. 13.**
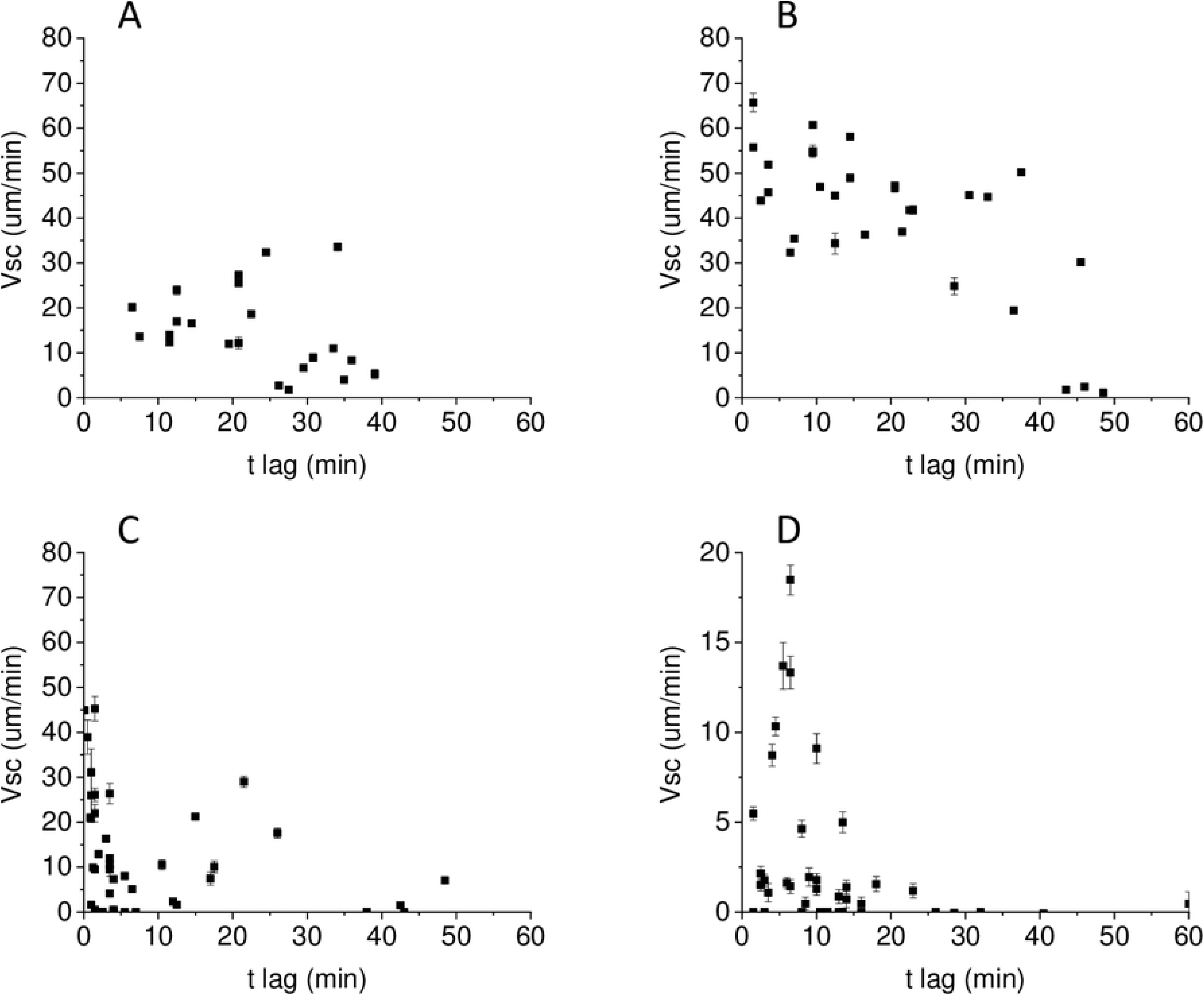
The rate of coagulation front propagation from centres of spontaneous clots and the lag time of clots appearance. (A) Data are represented for platelet MPs, (B) erythrocyte MPs, (C) endothelial MPs, and (D) monocyte MPs.

### MPs of different origins in coagulation propagation

The influence of MPs on coagulation propagation was evaluated by changes in the initial (Vi) and stationary (Vs) growth rates of clot growing from the activator in the Thrombodynamics test at different MPs concentrations (Fig. 14). Clot growth rates were calculated for MPs concentrations from 0 to those at which spontaneous clots were formed within 30 min, which prevented the correct calculation of the rates. Vi dependences on the concentrations of MPs of different origins tended to saturate with increasing concentration (Fig. 14 A, C). Vs dependences had the same tendency, but the influence of MPs was weaker and the saturation was less pronounced (Fig. 14 B, D). MMPs bearing the highest TF concentrations caused active spontaneous clotting at concentrations of 4-16·10^3^/µl, which did not allow the measurement of clot growth rate dependences on concentration in the same range as for other MPs types. At a concentration of 3·10^3^/µl, MMPs increased Vi by 5±6 µm/min and Vs by 2.4±2.2 µm/min. For THP MPs and EMPs bearing lower TF concentrations, the rate dependences on concentration could be measured to 50·10^3^/µl and 200·10^3^/µl, respectively. Significant differences in the rate dependences on the MPs concentrations of different origins were not observed in this range (Fig. 14 A, B). However, the Vi and Vs dependence on the ErMPs concentration tended to be smoother than the dependence on the MPs of other types at these concentrations. The measurement of rate dependences at higher concentrations was possible for PMPs and ErMPs only. The saturation levels of Vi were considerably different for these MPs types. The difference in Vs at high PMPs and ErMPs concentrations was even more pronounced (Fig. 14 A, C).

**Fig. 14.**
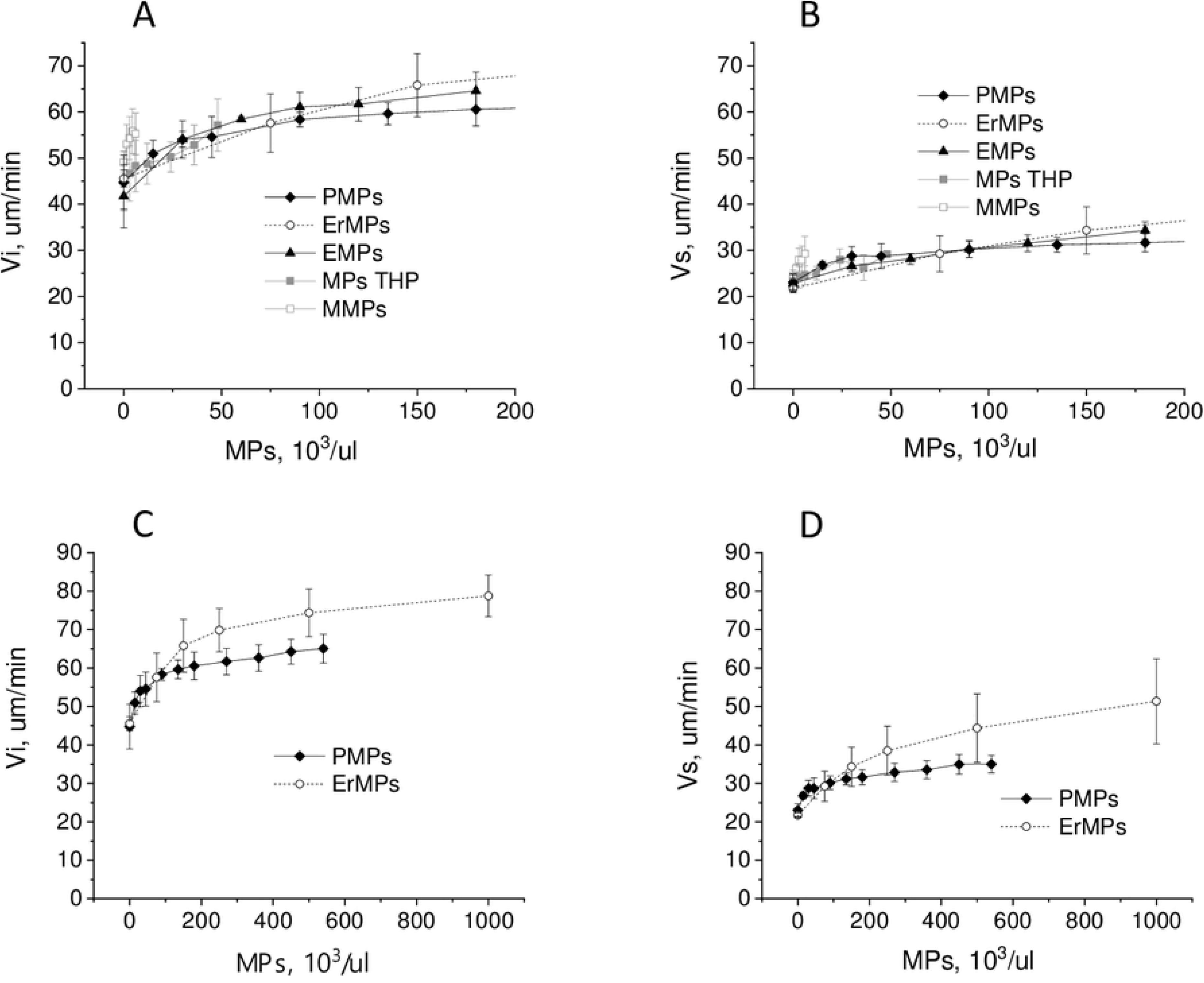
Influence of MPs of different origin on coagulation propagation. Mean ± sd dependence of the initial (A) and stationary (B) clot growth rates on platelet (PMPs) (n=10), erythrocyte (ErMPs) (n=7), endothelial (EMPs) (n=5), THP monocyte culture (THP MPs) (n=6) and monocyte (MMPs) (n=5) microparticles concentrations. (C), (D) The same dependences on a smaller scale.

It is possible that the difference between ErMPs and PMPs is explained by the different contents of phosphatidylserine (PS) because platelet membranes contain, according to various sources, from 6.7 to 12% PS [41–44], while erythrocyte membranes range from 13 to 16% [41,42,45]. To examine the effect of the PS content, we measured the dependence of clot growth rates on the concentration of artificial vesicles with a diameter of 100 nm, consisting of phosphatidylcholine (PC) and PS. The PS contents were 10%, 15%, and 20%. Based on the area occupied by one phospholipid molecule and the surface area of the vesicle, a concentration of 1 μM phospholipids corresponds to approximately 1.4·10^7^/μl of vesicles, which is approximately 20 times higher than the concentration of ErMPs causing spontaneous coagulation. Artificial vesicles did not induce spontaneous clotting at any of the tested concentrations. Both Vi and Vs reached saturation for artificial vesicles. The PS content increment from 10% to 20% led to a sharp increase in the slope of the initial linear part of the Vs dependence on the vesicle concentration from 0.24 µm/(min·µM) to 5.1 µm/(min·µM). Moreover, with an increase in PS content, the concentration at which the rates reached saturation decreased, but the saturation level did not change (Fig. 15). The maximal increase in Vs due to artificial vesicles was 13±3 µm/min. PMPs and ErMPs demonstrated a large deviation between donors. PMPs increased Vs by 11±4 µm/min on average and ErMPs by 23±8 µm/min at a concentration of 500·10^3^/µl. At a concentration of 1000·10^3^/µl, ErMPs increased Vs by 30±10 µm/min, and no saturation was observed. Therefore, both the type of dependence and the maximal effect indicate that the influence of ErMPs on Vs was not explained solely by the content of PS.

**Fig. 15.**
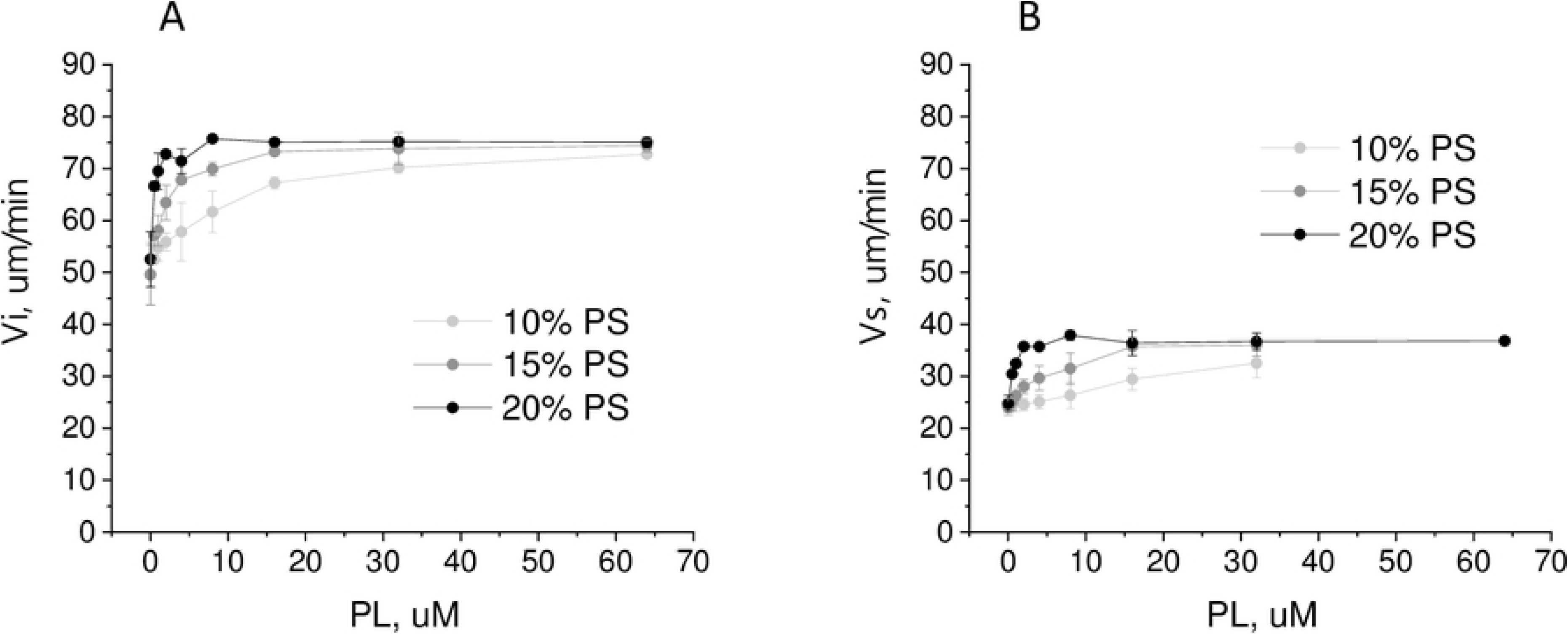
Influence of artificial vesicles on coagulation propagation. Mean ± sd dependence of the initial (A) and stationary (B) clot growth rates on the concentration of artificial phospholipid vesicles (PL) containing 10% (n=3), 15% (n=3) and 20% (n=2) PS.

## Discussion

It was previously shown that MPs in the plasma of patients cause the formation of spontaneous clots [46]. In the vast majority of cases, the clot growth rate from the activator in these samples was also increased. In this work, we investigated MPs of which origin has the strongest effect on the formation of spontaneous clots and the clot growth rate from the activator and, therefore, on coagulation activation and propagation. For this purpose, we tested MPs obtained in vitro from the main possible sources of MPs in blood in the Thrombodynamics test.

Our method of MPs counting with flow cytometry had some differences with the standard one: instead of counting annexin-positive MPs only, we took into account all the objects less than 1 µm and higher than the fluorescence threshold (the number of objects in stained buffer was subtracted). This was done because the percentage of annexin V-positive MPs (above the threshold of the negative control) significantly varied in MPs of different cellular origins: lower than 10%, 15-20%, approximately 30%, and up to 40% for ErMPs, PMPs, MMPs and THP MPs, and EMPs, respectively (see Materials and Methods). Variations in the percentage of annexin V-positive MPs could be at least partially explained by the difference in MPs size. Earlier, using the dynamic light scattering method, for the MPs sizing, we have shown in direct comparative studies that ErMPs have an average diameter of 200-250 nm, PMPs 350-400 nm, EMPs and THP MPs 400-500 nm, and EMPs 550-600 nm [29,30,47]. We presumed that annexin V-FITC binding to small MPs was too low to provide FITC signals above the threshold noise level, which is why we measured the different percentages of annexin V-positive events in samples of MPs of different origins with different average sizes.

It is difficult to evaluate the physiological range of MPs concentrations by comparing the absolute values of concentrations with literature data, since even when measuring concentrations using flow cytometry, depending on the measurement protocol and the cytometer used, normal MPs concentrations may vary from hundreds to 10^6^/μl [2,48–51]. Therefore, the physiological range of concentrations could be estimated only in relation to the normal MPs concentration measured in the same study. Our method of MPs counting is not applicable for measuring MPs concentrations in plasma. As a result, we can only assess the physiological range of concentrations indirectly. According to our data, the concentration of PS + MPs in the plasma is 53·10^3^/μl. The same concentration of PS + MPs is contained in approximately 300·10^3^/μl PMPs, 530·10^3^/μl ErMPs, 180·10^3^/μl MMPs and 130·10^3^/μl EMPs. Since ErMPs, EMPs, and MMPs normally make up less than 20% of the total MPs concentration [2], a concentration of 500·10^3^/μl for ErMPs and 200·10^3^/μl for EMPs and MMPs can serve as a rough estimate of the upper physiologically achievable concentrations.

The MPs activity in coagulation activation, estimated from the minimal concentration causing spontaneous clotting, was determined primarily by TF on the surface of the MPs, which corresponds to the results from comparing the same MPs in the recalcification test in our previous works [29–31] and in the thrombin generation test in [24–26]. Based on the fact that MMPs, such as THP MPs, have ∼ 30% PS+MPs, the ratio of PS+MMPs leading to spontaneous clotting to the normal concentration of PS + MPs in plasma will be approximately 4%. Such a concentration seems quite achievable in vivo, especially locally, for example, for inflammatory vascular diseases, monocytes and endothelium activation.

The ratio of the number of coagulation centres to the number of MPs in a chamber is on the order of one clot per 10^3^-10^7^ MPs. This led to the assumption that coagulation centres may be caused not by MPs themselves but by some larger residual fragments of cells or MPs aggregates. In addition to the fact that the ratio of the number of clots to the number of MPs is greater for TF-bearing MMPs and EMPs, coagulation begins over the whole volume of the chamber, not just from the individual centres, when certain concentrations of MMPs and EMPs are reached. This is probably due to the participation of a much larger fraction of TF+MPs than 1/10^3^ in coagulation activation due to the effect described by Kastrup et al., when the convergence of several centres with individual subthreshold activation leads to overcoming the activation threshold [52].

In the range up to 200·10^3^/μl, there were no significant differences in the effect of PMPs, ErMPs, and EMPs on the clot growth rate. Perhaps we could not identify it due to deviations between the MPs samples isolated from the cells of different donors and between the donor plasmas in which the titrations were performed. However, in view of the fact that the differences in the effects of MPs on the rates were weak and the protein composition in MPs originating from different cells is also different, the effect of MPs on rates is probably determined by the lipid composition of membranes to a larger extent than by the protein composition. Rather inactive at low concentrations, ErMPs at 500·10^3^/μl increased the clot growth rate to a significantly higher level in comparison with 500·10^3^/μl PMPs. The growth rate of spontaneous clots induced by ErMPs also significantly exceeds the growth rate of clots induced by PMPs, EMPs and MMPs. The high ErMPs activity in coagulation propagation is consistent with the data of van der Meijden, where the same conditional PS activity, which was measured by the prothrombinase activity on MPs samples, corresponded to a 3-fold higher activity of PMPs compared with that of ErMPs [26]. ErMPs activity is not explained by PS only because artificial vesicles lead to clot growth rates that are significantly lower than those of ErMPs at concentrations higher than 500·10^3^/μl, and an increase in the PS content in artificial vesicles did not change the saturation level. Other components of phospholipid composition may determine ErMPs activity. Notably, ErMP concentrations could be underestimated due to their small size. Thus, the difference between ErMPs and PMPs can partially be accounted for by higher ErMPs concentrations, and the attainability of 500·10^3^/μl ErMPs in physiological conditions is questionable.

A significant distinction of TF-bearing MPs is that there are concentrations at which MPs are able to induce some separate clots, but the growth of these clots stops in the first 10-15 min of a test. Previously, Oliver et al. concluded, by means of the thrombin generation test, that TF-bearing MPs participate in coagulation activation but not in coagulation propagation [53]. It is likely that clot growth stops when subthreshold activation leads to thrombin impulses that are rapidly inhibited by plasma inhibitors, and the lipid surface may not be enough to support coagulation propagation because these concentrations are quite low. Another reason for the growth termination of EMPs and MMPs induced clots could be thrombomodulin on the surfaces of endothelium and monocytes [54, 55].

Summarizing the results, we can say that MPs derived from different cells play a qualitatively different role in coagulation activation and propagation: TF+ MMPs have a strong activating ability and have a very weak effect on coagulation propagation; on the contrary, contact activation from PMPs and ErMPs in normal plasma is weak, and these MPs, firstly, contribute coagulation propagation. Although an increased concentration of MPs is usually regarded as a risk of thrombosis, MPs that have weak activating capacity but support coagulation propagation in some cases can play a positive role, for example, by reducing blood loss during surgery or mitigating the clinical manifestations of haemophilia [56]. Endothelial MPs, although they have an intermediate activity, are able to make a significant contribution to both the activation and distribution of coagulation.

## Acknowledgments

We thank Dr. Panteleev MA (Center for Theoretical Problems of Physicochemical Pharmacology, Moscow, Russia) for valuable discussions.

The study was supported by the Ministry of Science and Higher Education of the Russian Federation (project АААА-А18-118012390250-0) and by the Russian Foundation for Basic Research together with National Center for Scientific Research of France (grant 19-51-15004 to Ataullakhanov F.I.).

## Supplementary materials

S1 Movie. Video of clots growing from activator and spontaneous clots in MP-depleted plasma supplemented with MPs of different origins in different concentrations.

Fig. S1. Dependence of the time of the first 5 spontaneous clots appearances on time. Data are represented for (A) platelet MPs, (B) erythrocyte MPs, (C) endothelial MPs, and (D) monocyte MPs. Dots correspond to individual tests, the mean values of t_N=5_ at different concentrations are connected with lines, and symbols of different types and colours correspond to different MPs samples.

Fig. S2. Comparison of the maximal light scattering intensity in the centres of spontaneous clots and of clots growing from activator. The maximal light scattering intensity dependence on the concentration of (A) platelet MPs, (B) erythrocyte MPs, (C) endothelial MPs, and (D) monocyte MPs. The maximal light scattering intensity in the centres of spontaneous clots is denoted with opened symbols, and that of clots growing from activator is denoted with filled symbols.

Fig. S3. Concentration dependence of the maximal light scattering intensity in the centres of spontaneous clots. Data represent (A) platelet MPs, (B) erythrocyte MPs, (C) endothelial MPs, and (D) monocyte MPs. Dots correspond to individual tests, the mean values of VI at different concentrations are connected with lines, and symbols of different types and colours correspond to different MP samples.

Fig. S4. Concentration dependence of the rate of coagulation front propagation from centres of spontaneous clots. Data represent (A) platelet MPs, (B) erythrocyte MPs, (C) endothelial MPs, and (D) monocyte MPs. Dots correspond to individual tests, the mean values of VI at different concentrations are connected with lines, and symbols of different types and colours correspond to different MP samples.

